# DLITE uses cell-cell interface movement to better infer cell-cell tensions

**DOI:** 10.1101/541144

**Authors:** R. Vasan, M.M. Maleckar, C.D. Williams, P. Rangamani

## Abstract

Cell shapes and connectivities evolve over time as colony shapes change or embryos develop. Shapes of intercellular interfaces are closely coupled with the forces resulting from actomyosin interactions, membrane tension, or cell-cell adhesion. While it is possible to computationally infer cell-cell forces from a mechanical model of collective cell behavior, doing so for temporally evolving forces in a manner that is robust to digitization difficulties is challenging. Here, we introduce a method for Dynamic Local Intercellular Tension Estimation (DLITE) that infers such temporal force evolutions with less sensitivity to digitization ambiguities or errors. This method builds upon prior work on single time points (CellFIT). We validate our method using synthetic geometries. DLITE’s inferred cell colony tension evolutions correlate better with ground truth for these synthetic geometries than tension values inferred from methods that consider each time point in isolation. We introduce cell connectivity errors, angle estimate errors, connection mislocalization, and connection topological changes to synthetic data and show that DLITE has reduced sensitivity to these conditions. Finally, we apply DLITE to time series of human induced pluripotent stem (hIPS) cell colonies with endogenously expressed GFP-tagged ZO-1. We find major topological changes in cell connectivity, e.g. mitosis, can result in an increase in tension. This supports a correlation between the dynamics of cell-cell forces and colony rearrangement.

**Significance statement:** Cell-cell tensions play an important role in the dynamics of tissue morphogenesis. Mathematical modeling tools have helped understand the role of cell-substrate and cell-cell adhesion in tissue organization. In particular, recent modeling studies have shown that an inferential approach without a constitutive equation can estimate distribution of tensions in a single image of a cell monolayer (CellFIT). Here, we include the dynamics of monolayer morphogenesis in the estimation of cell-cell tensions. Such a formulation, termed DLITE, performs better across time in both synthetic geometries and time series of hIPS cell colonies with endogenously expressed GFP-tagged-ZO-1. We also show that DLITE is robust to digitization ambiguities during segmentation. Such a method can shed some light on physical mechanisms that drive morphogenesis.

## Introduction

Cell shape, forces, and function are closely related [1, 2, 3, 4]. Cell shape affects the organization and transmission of cytoskeletal forces and the structures that create them [5, 6, 7]. Cell membrane tension partially governs processes from intracellular endocytic bud morphology during trafficking [8] to tissue level remodeling events such as wound healing [9], development [10], expansion [11], migration [12, 13] and cancer invasion [14]. Mechanical rearrangement occurs as cells transmit these forces across the membrane [15] and cell-cell adhesion complexes such as adherens and tight junctions [16, 17]. These apical cortical complexes [18] depend on the activity of the actomyosin cytoskeleton [19, 20]. The mechanotransduction of intercellular forces can alter and regulate biochemical signalling pathways [21, 22] with force and deformation at a particular time point partially regulating future force and deformation.

Force-mediated collective behaviours are crucial for the dynamics of tissue reshaping. This is commonly evidenced by apoptosis in cell cultures or by the intercalation and extrusion of cells during development [23, 24]. We can examine the role of tension in tissue remodeling using direct force measurement techniques such as atomic force microscopy (AFM) or micro-pipette aspiration [25, 26, 27]. These direct measurement techniques offer precise force characterization in cells and tissues but perturb the actomyosin network [28, 29]. As a result, these methods can alter the force responses of the system at subsequent time points. Alternative optical measurement techniques that use Förster Resonance Energy Transfer (FRET) tension probes or traction force microscopy (TFM) can assay force [30, 31, 32] without the mechanical disruption associated with direct measurements [33]. These optical approaches can be applied across extended periods but, like the prior physical techniques, they can be difficult to implement in a high-throughput context. Each of these measurement techniques has its distinct set of advantages, depending on the biological problems being studied. Complementary to these approaches, force inference from the geometry of the cell boundary can allow for the estimation of normalized tensions solely from images of labeled confluent cells without further condition requirements.

Intercellular forces can be inferred at cell-cell interfaces using a mechanical model predicated on the assumption that forces are balanced where multiple cell-cell interfaces meet [34, 35, 36]. These mechanical models cover a range of complexities, assumptions, and use cases [37]. Here we deal with the subset devoted to the representation of tensions in a two-dimensional plane of curved edges digitized from the apical interfaces of a confluent cell colony (Fig. 1A,B). We build upon prior representations of this system, notably that used in the Cellular Force Inference Toolkit (CellFIT) [34], and develop an alternate problem formulation that treats tension estimation as a temporally evolving problem, borrowing information from prior time points to increase model prediction stability and boost resistance to ambiguities or errors that arise during the digitization process (Fig. 1C). This provides a non-disruptive means to infer intercellular forces in time-lapse imaging of cell colonies. We term this technique Dynamic Local Intercellular Tension Estimation, or DLITE. Here, we validate DLITE against a range of synthetic data, for which known tension ground truths are available, and use it to predict tensions in time series of human induced pluripotent stem (hIPS) cell lines with the endogenously GFP-labeled tight junction protein ZO-1.

**Figure 1:**
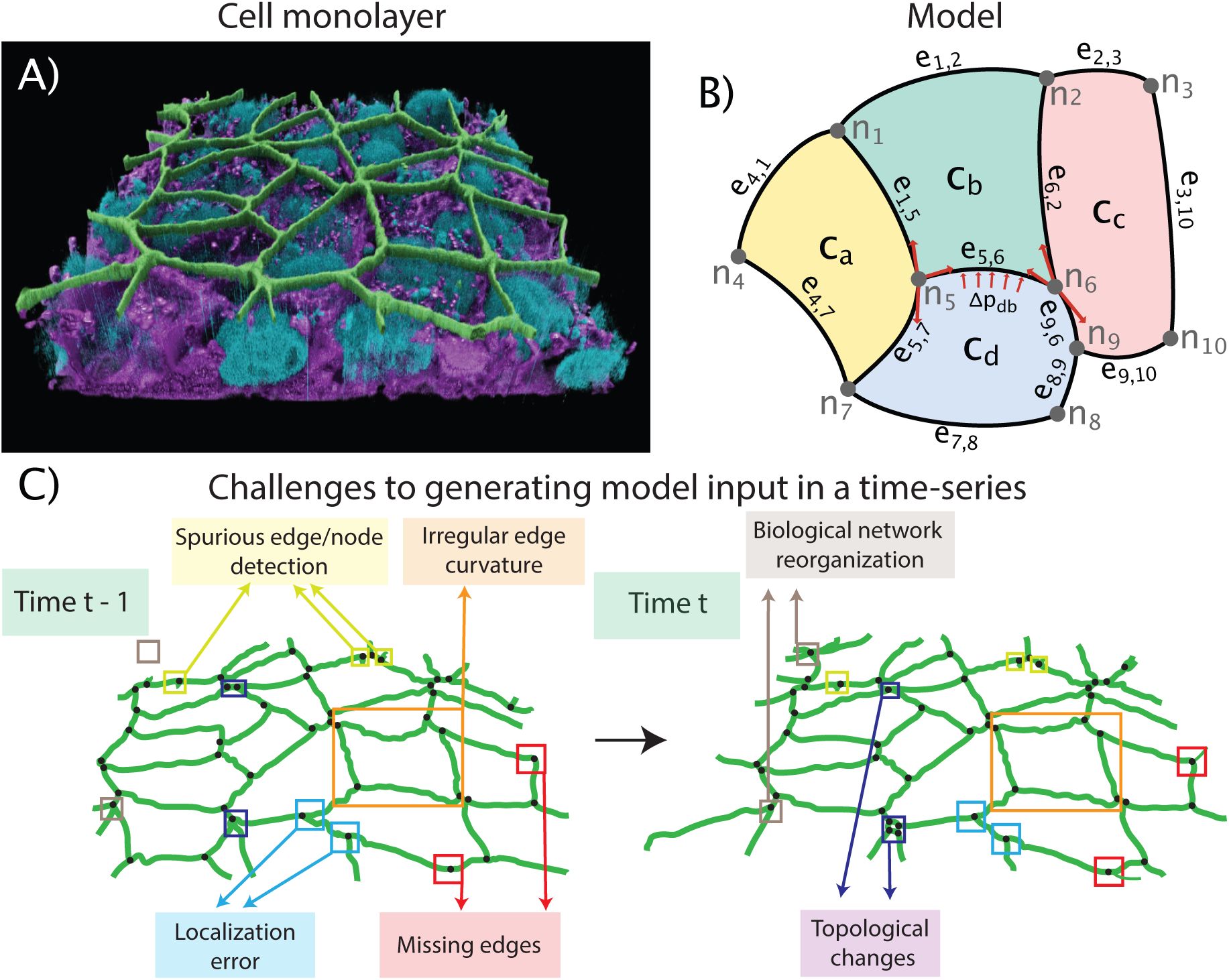
3D cell view of tight junction location, how this is represented in the model, and the challenges in doing so. (A) 3D view of tight junctions in human induced pluripotent stem (hIPS) cells from the Allen Cell Explorer (Green - Tight junctions, Purple - Membrane, Blue - Nucleus). We infer cell-shape and edge shape from tight junctions as they localize to the tension bearing apical surface of epithelial-like tissues. (B) Schematic of cell-interface representation used in DLITE and CellFIT force-inference techniques [34]. A colony is represented as a set of nodes (*n*), edges (*e*) and cells (*c*). Edges are directional. Tension balance occurs at each node (red arrows at n_5_ and n_6_). Pressure difference (∆*p*_*d,b*_) across a junction is estimated using Laplace’s law (red arrows at e_5,6_). (C) Ambiguities in image segmentation introduce challenges to successful tension inference. Time t - 1 shows single time point challenges like spurious edge/node detection, irregular edge curvature, node location errors and incomplete segmentation. Time t shows time lapse challenges like biological network reorganization and topological changes.

## Methods

### Assumptions

We employed a curvilinear description of a tissue by defining a colony as a directed planar graph comprising cells (*c*), edges (*e*) and nodes (*n*) (Fig. 1B, for more details see SOM and [38, 34]). Forces exerted by the actomyosin cortex result in tangential stresses in the form of tension (*t*) along an edge. Cells resist deformation by means of a normal stress exerted as pressure (*p*) inside every cell. Along each edge, we assumed that the interfacial tensions are constant and that the intracellular pressures are uniform within a cell. At the length scale of the whole cell, we ignored membrane bending and assumed that edge tensions and cell pressures exclusively govern cell shape. We treated viscous forces as negligible and therefore assumed that the colony shape is quasi-static, i.e. at each time point the colony is in mechanical equilibrium.

### Governing equations and system specification

A general force balance at every node in a colony can be written as

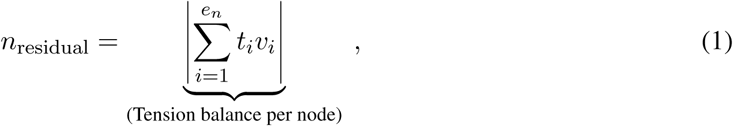

where *n* is a node, *t*, and *v* represent the tension/tangential stress and local tangent unit vector of an edge connected to node *n* respectively, *e*_*n*_ is the number of edges connected to node *n* and *n*_residual_ is the magnitude of the resultant tension vector coming into a node (ideally 0). This notation is shown in Fig. 1B. This equation applies when employing a curvilinear description of tissue and applies to a node that is both connected to at least three edges and is in mechanical equilibrium [34]. The pressure difference between adjacent cells can be estimated using Laplace’s law as

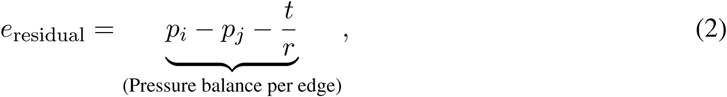

where *e* is an edge, *p*_*i*_ and *p*_*j*_ are the pressure of adjacent cells *i* and *j* respectively and *e*_residual_ is the residual error from the pressure balance. Here, *t* and *r* represent the tension and radius of the interfacial edge *e*. The system of equations for tension and pressure are generally overdetermined; there is no unique solution to this system [34]. Therefore, we can only infer the relative distribution of tensions from the shape of the edges and not the absolute values.

To compute the dynamics of cell-cell forces, we reformulated the tension balance (Eq. 1) as a local optimization problem defined as

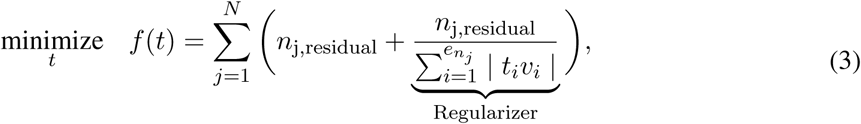

where *n*_*j*_ and *e*_*n*_*j*__ represent the j^*th*^ node and the number of edges connected to node *n*_*j*_, and *N* is the total number of nodes in the colony. Here, *n*_*j*,residual_ is the tension residual at a given node (Eq. 1) and the regularizer is the magnitude of the tension residual divided by the sum of the magnitude of the tension vectors acting on that node. Since tension cannot be negative, we set a lower tension bound of zero. In Eq. 3, the regularized term ensures that the system of equations does not converge to the globally trivial solution (tension = 0 along all edges) [39]. Pressure in each cell was computed using

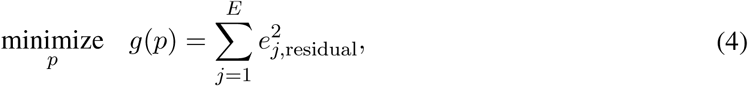

where E is the total number of edges in the colony and e_*j*_ is the residual error from the pressure balance at the j^*th*^ edge. Tension and pressure solutions were normalized to an average of 1 and 0 respectively, similar to prior work [34, 36, 35]. In contrast to previous methods, DLITE uses the values of tension at each edge and pressure in each cell from the previous time point as an initial guess for the current timepoint. This mode of time-stepping in the optimization procedure enables us to use information from previous time points to predict the values of tension and pressure at the current time point and forms the basis of DLITE’s improved performance across time-series.

### Tracking nodes and edges

An essential distinguishing characteristic of DLITE is the ability to provide an initial guess for each edge tension and each cell pressure, allowing us to incorporate a time history of cell-cell forces. However, this requires node, edge, and cell tracking over time. To implement tracking we first assign labels to nodes, edges, and cells at the initial time point. Then, nodes are tracked by assigning the same label to the closest node at the next time point. Edges are tracked by comparing edge angles connected to nodes with the same label and cells are tracked by matching cell centroid locations across time. Our model optimization pipeline was implemented using Scipy’s unconstrained optimization algorithm ‘Limited-memory Broyden–Fletcher–Goldfarb–Shanno (L-BFGS)’ [40]. The global optimization technique ‘Basinhopping’ was used to seek a global minimum solution at the first time point [41].

### Geometries for model validation

Validation of DLITE requires the generation of dynamic 2D geometries with curvilinear edges whose cortical tensions are known. Many standard mathematical models describe the modification of cell-shape via applied forces that are either explicitly or implicitly specified. Such models include cellular Potts models [42, 43], Vertex models [44, 45] and cell-level finite element models [46, 47, 48]. Implicit models define an energy function relating the variation of tension and other properties in a 2D monolayer to cell shape. The gradient of this energy function leads to the movement of each vertex. Here, we employ an implicit model using the energy minimzation framework Surface Evolver [49], which is designed to model soap films. The energy function (*W*) was defined as

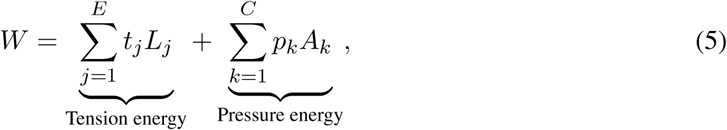

where *t*_*j*_, *L*_*j*_ are the tension and length of the j^*th*^ edge and *p*_*k*_, *A*_*k*_ are the pressure and area of the k^*th*^ cell respectively. *E* and *C* are the total number of edges and cells in the colony (see SOM for details). Here, the tension energy represents a net energy contribution due to adhesion forces that stabilize a cell-cell interface and actomyosin cortical tensions that shorten cell-cell contacts. Pressure was enforced as a Lagrange multiplier for an area constraint. Cell boundaries were free to move along the surface. Such a model outputs a minimum energy configuration through gradient descent, providing ground truth tensions to which we compare inference model outputs. While the model utilized here describes a monolayer as a 2D surface embedded in 3D space, it is possible to extend this work to 3D, covering the complex 3D structure present in many systems [36].

### Sources of error due to digitization

Transforming single or multi-channel z-stacks of cell colonies into a connected network suitable for tension inference requires: *(i.)* Image pre-processing to produce a binary or otherwise simplified representation, *(ii.)* Skeletonization, creating a network of 0-width lines connecting nodes at junction points and *(iii.)* Post-processing of the skeletonized representation. Inherent ambiguities in this process introduce several challenges to successful tension inference. Some of these challenges, such as incorrectly detecting an edge, occur in single frames (Fig. 1C, Time t-1) while others, such as edge tracking across biological network reorganization, are only present in time series data (Fig. 1C, Time t). These challenges tend to occur more frequently as digitization is increasingly automated, creating a trade-off between data reliability and throughput.

## Results

DLITE is built to use the tension at a specific cell-cell junction at a given time point as an initial guess to calculate tensions at the next time point. This logical progression then allows us to infer forces over time and test the strength of the inference method by correlation to ground truth values for synthetic geometries. We demonstrate robustness and sensitivity of DLITE by validating it against ground-truth tensions for multiple synthetic geometries, multiple tension perturbations within a colony, connectivity ambiguities at single or multiple time points, curve-fit errors, node location errors, and topological changes like the shrinkage of cell-cell contacts. At each point, we compare predictions to those produced by the state of the art CellFIT technique. We then apply DLITE to movies of skeletonizations of endogeneously tagged tight junction ZO-1 (*zonulae occludentes*-1) in an hIPS cell line and demonstrate improved tension stability in the inference of cell-cell forces during colony dynamics.

### Validation of DLITE as a dynamic tension-inference tool

We validated DLITE in three steps. First, we compared edge tension and cell pressure solutions obtained using our implementation of the CellFIT algorithm and DLITE to ground truth values in synthetic geometries made available via the current version of CellFIT called ZAZU (Fig. S1). We re-implemented the CellFIT algorithm in Python because the source code of ZAZU is not publicly available. Both our re-implementation of CellFIT (Fig. S1B) and DLITE (Fig. S1C) perform identically with respect to the ground truth for single frames (Fig. S1A), with an average error of *∼* 0.02 (Fig. S1D).

Second, to generate a time-series of synthetic geometries, we simulated colonies that deformed smoothly across time using Surface Evolver [49]. Initial geometries were created from random Voronoi tessellations followed by Lloyd relaxation [50]. In order to generate a time-series, we performed multiple Evolver simulations where the tension of a few randomly selected edges were either increased or decreased between time points (Fig. 2A). Average edge tension at each time point was normalized to 1. We then stripped all tension and pressure information from the resulting shapes and used these shapes as input to our method. Using DLITE and CellFIT separately, we inferred colony forces and compared the two approaches (Fig. 2B, D). Initially, both methods performed identically, but began to show divergence after 3 frames. Importantly, we observed that the values of tension predicted using our method remain closer together over time and are better correlated (r = 0.94 (DLITE) vs r = 0.75 (CellFIT)) to the ground truth (Fig. 2 B, D). Further, we observed that the variation in tension defined by the change in edge tension over time denoted as Δtension, using DLITE also correlates better to the ground truth change in edge tension (Fig. 2C). The improved performance of DLITE at later time points (Fig. S2B) results from DLITE use of information from prior time points to improve tension predictions in the presence of large curve-fit residuals (Fig. S2A), thereby reducing sensitivity to curve-fitting errors. In the absence of an informed prior, i.e. when we use random initial guesses sampled from a random Gaussian distribution at every time point, we observed poor performance of DLITE, with correlations ranging from 0.75 to 0.89.

**Figure 2:**
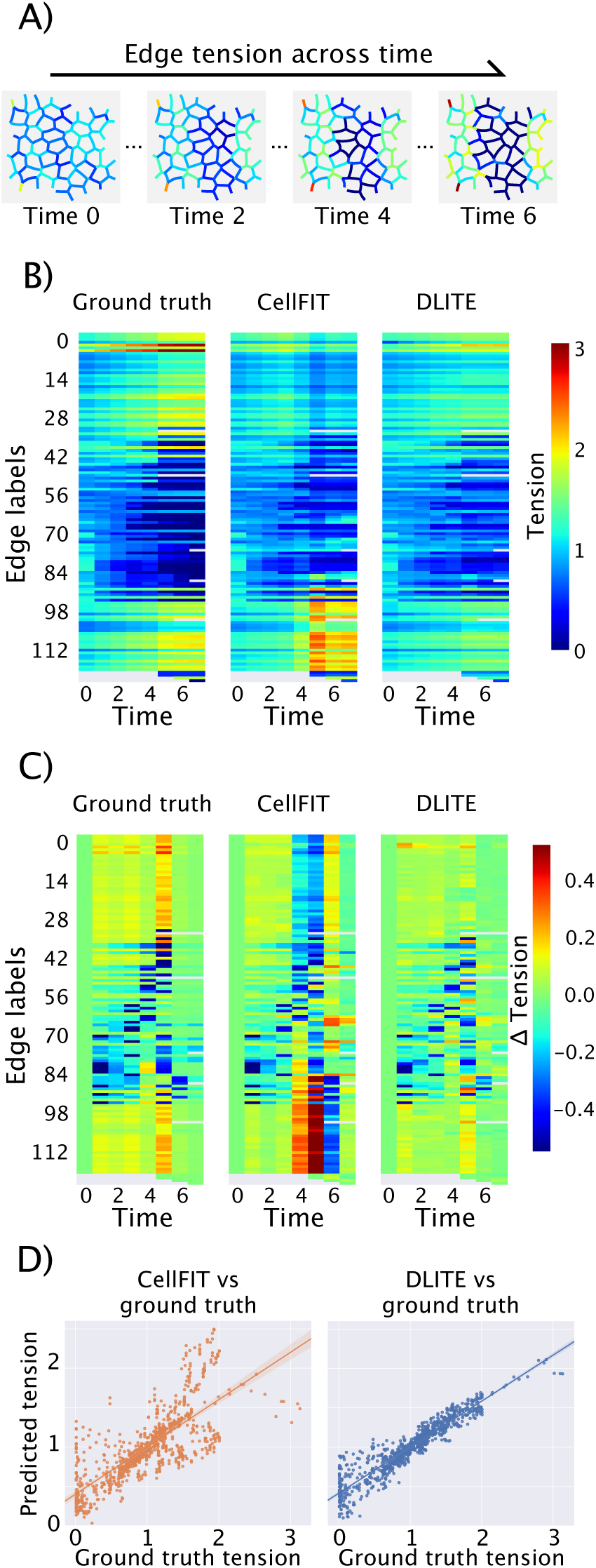
Comparison of DLITE and CellFIT force-inference techniques for digitized time series. Synthetic colonies were generated from random Voronoi tessellations and morphed to minimum energy configurations (Eq. 5) using Surface Evolver [49]. A random set of edges within the colony were perturbed by decreasing or increasing their tensions, resulting in a new colony structure; repeating this process produced a time-series of colony rearrangement. (A) Time-series of a synthetic colony showing the decrease in tension of 70 edges in the middle of the colony and the increase in tension of 40 edges along the boundary. (B) Heatmap of dynamic edge tensions for ground truth, CellFIT, and DLITE. (C) Heatmap of dynamic change (derivative of tension) in edge tensions for ground truth, CellFIT, and DLITE. (D) A comparison of inferred vs ground truth tensions for CellFIT (r = 0.75) and DLITE (r = 0.94). Here, r is the Pearson’s correlation coefficient.

Third, to ensure robustness of the performance of DLITE, we tested multiple tension perturbations via different combinations of increasing and decreasing edge tension in the same geometry (Fig. S4) and similar perturbations in other randomly generated geometries (Fig. S5). In all cases, we observed equivalent or better correlation of both the tension and change in edge tension with the ground truth using DLITE as compared to CellFIT.

### DLITE is robust to digitization ambiguities

The input to a force-inference model is a map of colony shape as a series of curved edges and the nodes where edges join (Fig. 1B). Segmentation transforms image data into information about the isolated geometric structures [51, 52]. Subsequently, skeletonization methods extract lines that characterize the topology and connectivity of the tension bearing network in the colony. Ambiguities or errors in this mapping present challenges to force inference techniques that rely on precise colony connectivity and edge tracing [34, 35, 36]. Some of these conditions are shown in Fig. 1C. New methods have improved the quality and repeatability of predicted network topology and connectivity; both deep learning models and traditional computer vision techniques have made significant advances in 2D/ 3D biological segmentations [53, 54, 55, 56]. Despite these advances, current skeletonization methods continue to require semi-manual post-processing because of the ambiguities present in the structures during imaging and errors resulting from the image capturing modalities. This semi-manual cleanup becomes increasingly impractical for larger colonies and time series. As a result, we require force-inference techniques robust to errors in mapping. As we demonstrate below, our inference method has increased robustness to multiple edge/node mapping errors. Therefore, our method decreases the number of manual corrections required, and increases the tractability for inferring forces. Here we evaluate the effects of edge/node mapping errors on force-inference in a single image and in a time-series.

We first analyzed, at a single time point, the effect of a missing intersection between two edges, a commonly occurring connectivity error. As before, we generated a synthetic colony image to initialize the system with known edge tensions. Fig. 3A shows a random Voronoi tessellation generated using Surface Evolver where 50 edges (out of 330 total) have larger values of tension than others. The ground truth tensions were the inputs given to Surface Evolver (Fig. 3A). A single edge was deliberately traced incorrectly to introduce a connectivity error (Fig. 3A, *inset*). This error resulted in the loss of a triple junction and loss of cellular integrity. Since the node of interest is now connected to two edges instead of three, we can no longer conduct a tension balance at that location. Such an ill-posed problem results in a singular tension matrix G_*γ*_ (Eq. 6), implying that CellFIT is unable to infer a correct tension distribution (Fig. 3A&B). However, the use of a regularizer in DLITE (Eq. 3) reduces the effect of local tension errors on the global data set. As a result, we find that at a given time point, DLITE is able to provide a good estimate of the tension of the neighboring edges, even in the presence of connectivity errors (Fig. 3C, see also Figs. S6, S7).

**Figure 3:**
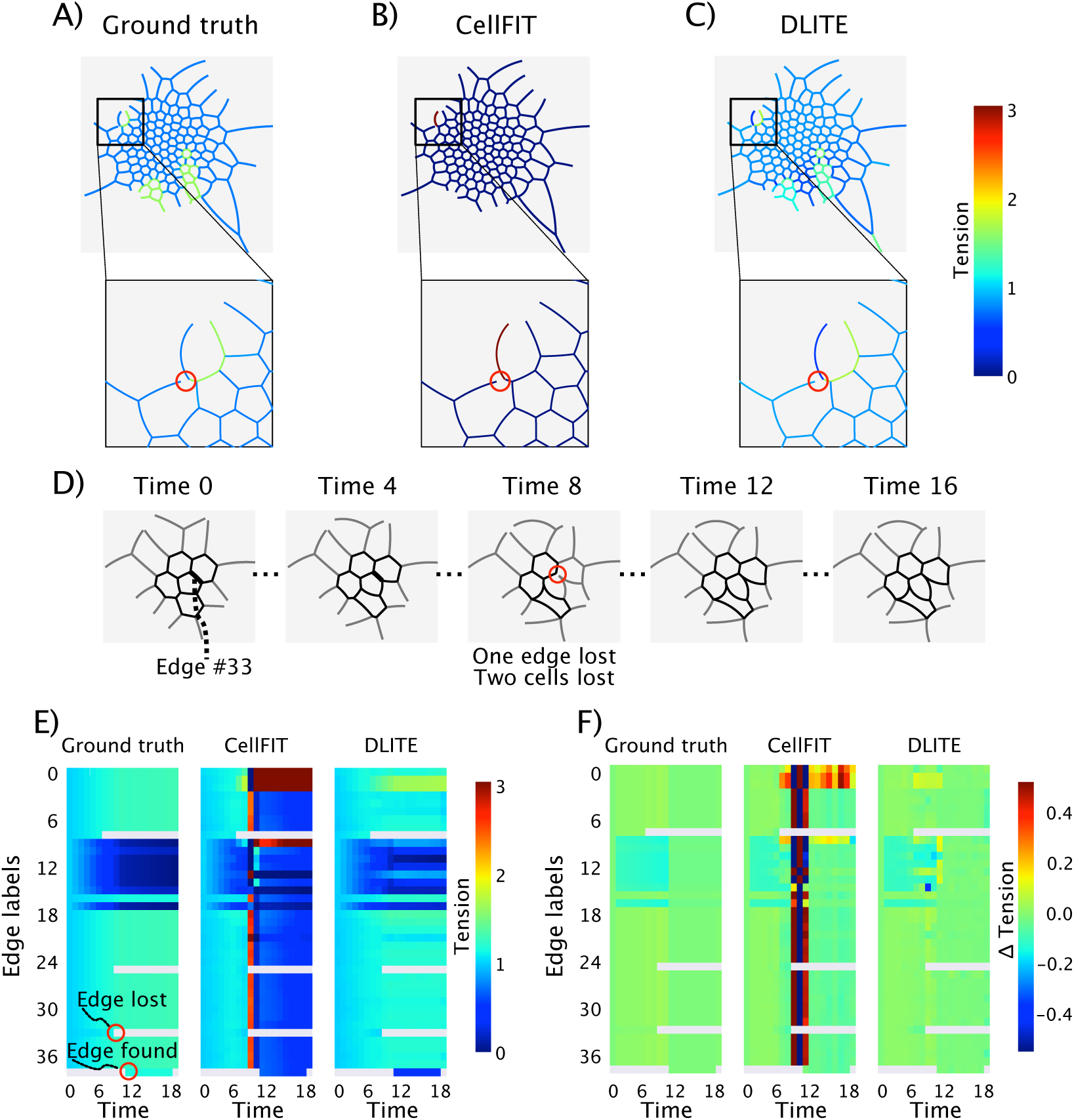
Reduced sensitivity to connectivity errors in DLITE. (A) Ground truth tensions for a synthetic geometry containing 330 edges generated used Surface Evolver with a single edge connectivity error (circled in red). (B) Edge tensions computed using CellFIT for the geometry in (A). (C) Edge tensions computed using DLITE for the geometry in (A). (D) Time-series of a synthetic geometry containing 37 edges generated using Surface Evolver with a single edge connectivity error at time 8 (circled in red). This edge is found again in time step 10 (representing a transient encoding error) but treated as a new edge. (E) Heatmap of dynamic edge tensions for ground truth, CellFIT (r = 0.14) and DLITE (r = 0.87) for the time-series in (D). (F) Heatmap of dynamic change (derivative of tension) in edge tensions for ground truth, CellFIT, and DLITE for the time-series in (D).

A commonly occurring digitization challenge results from poor estimation of edge curvature due to incorrect values of between-edge angles at a particular node. Errors in curve-fitting can lead to poor tension residuals (Eq. 1) or large condition numbers of tension matrices (Eq. 6), which is defined as the ratio of the largest to smallest singular values in the SVD (Singular Value Decomposition) of the given tension matrix. Subsequently, this leads to poor inference of tension (Fig. 2). This is especially problematic when cell-cell junctions are distinctly non-circular, as they commonly are. To simulate this, we generated a time-series of synthetic geometries using Surface Evolver such that later time points are distinctly non-circular (Fig. S3A). The large curve-fit residuals at later time points (Fig. S3B) lead to ill-conditioned tension matrices and errors in tension inference (Fig. S3C). However, DLITE uses tension information from prior time points to retain the distribution of tensions and is not poorly scaled by these curve-fit errors (Fig. S3C).

Another major digitization challenge for force-inference models is the accurate determination of node locations. Localization errors in node coordinates also have the downstream impact of changing connected edge curvatures (Fig. 5B). We simulated this type of error by adding levels of Gaussian noise to nodes in a synthetic colony (Fig. 5A, red nodes). Noise levels 1 (Fig. 5C, F), 2 (Fig. 5D, G) and 3 (Fig. 5E, H) refer to the Gaussian noise terms with a mean of 0 and standard deviations of 0.1, 0.5, and 1 respectively. Red node coordinates are (480.95, 525.7), (487.76, 536.94), (498.63, 522.1), (524.25, 503.43), (535.62, 515.97), arranged from left to right. In all cases, we observed equivalent (Noise level 1) or improved performance (Noise levels 2, 3) when using DLITE compared to CellFIT. Thus, DLITE offers improved quality of tension inference in the presence of ambiguities in node location.

Finally, we considered a class of mapping challenges that are unique to time-series data – identification of edges from one frame to the next. Misidentification of edges frequently occurs when an edge is lost for a single frame, severing the edge’s connection to its prior label. Fig. 3D shows a time-series with a missing edge (edge label 33) and two missing cells at time point 8. The missing edge leads to the loss of a triple junction, and consequently a singular tension matrix (Fig. 3E, CellFIT). For tracking purposes, missing an edge also means that the edge that was being tracked up to that point no longer exists, and therefore a new edge label is assigned. Since edges with new labels do not have an initial tension guess from an identical label at prior time points, these edges are given an initial guess for the value of tension equal to the average initial guess of all edges connected to that edge. By using such a scheme, DLITE can predict tension and Δtension (change in edge tension of an edge label between adjacent time points) that correlates well with the ground truth (Fig. 3E, F). Thus, in both images and movies of colonies, we find that use of information from the neighbouring region allows DLITE to handle digitization ambiguities and errors better and robustly predict the distribution of cell-cell forces.

### DLITE is robust to topological changes

Network topology or the structure of edges and vertices often display changes in time-series data. In Fig. 3D, for example, there are two topological changes at time points 6 and 8 (edge labels 8 and 25 respectively) that result in differences between CellFIT and DLITE (Fig. 3E, F), with DLITE showing better correlation to the ground truth. Handling of a time-ordered network requires the tracking of nodes, edges, and cells over time. This can be done in 2 ways – if, for example, a single edge ceases to exist at a certain time point, we can choose to either keep that edge label and assume that it has temporarily left the field of view or assign new edge labels ensuring that the lost edge label ceases to exist [57]. Here, we choose the second option in order to condition the network based only on the immediately prior time point. Thus, an edge label that could not be tracked after a time point no longer exists and is assigned a new label. These rules were applied to nodes and cells as well.

If the observed topologies of a cellular network are constantly changing, how then does it affect inferred cell-cell forces? To study the effect of topological changes, we take advantage of the fact that decreasing the tension of two edges in a triple junction results in a decrease of the length of the third connected edge in Surface Evolver. Fig. 4A shows an example time series where edge label 15 disappears at time point 18. This single topological change leads to an ill-conditioned tension matrix (Eq. 6, Fig. 4B, C). However, DLITE retains the correct distribution of tensions at time point 18 (see also Fig. 3E, edge label 25 at time point 8 and Fig. S8A) using the initial guess from prior time points. While this specific network structure led to an ill-conditioned tension matrix after a single edge loss, this is not always the case. If the tension matrix is well-conditioned after the topological change (Fig. 3E, edge label 8 at time point 6 and Fig. S8B), then CellFIT retains a good solution quality. However, Δtension is less smooth, and both the tension and Δtension still correlate better to the ground truth while using DLITE (Fig. 3E,F and Fig. S8B). Thus, force inference in dynamic network topologies benefits from incorporating a temporal history of cell-cell forces.

**Figure 4:**
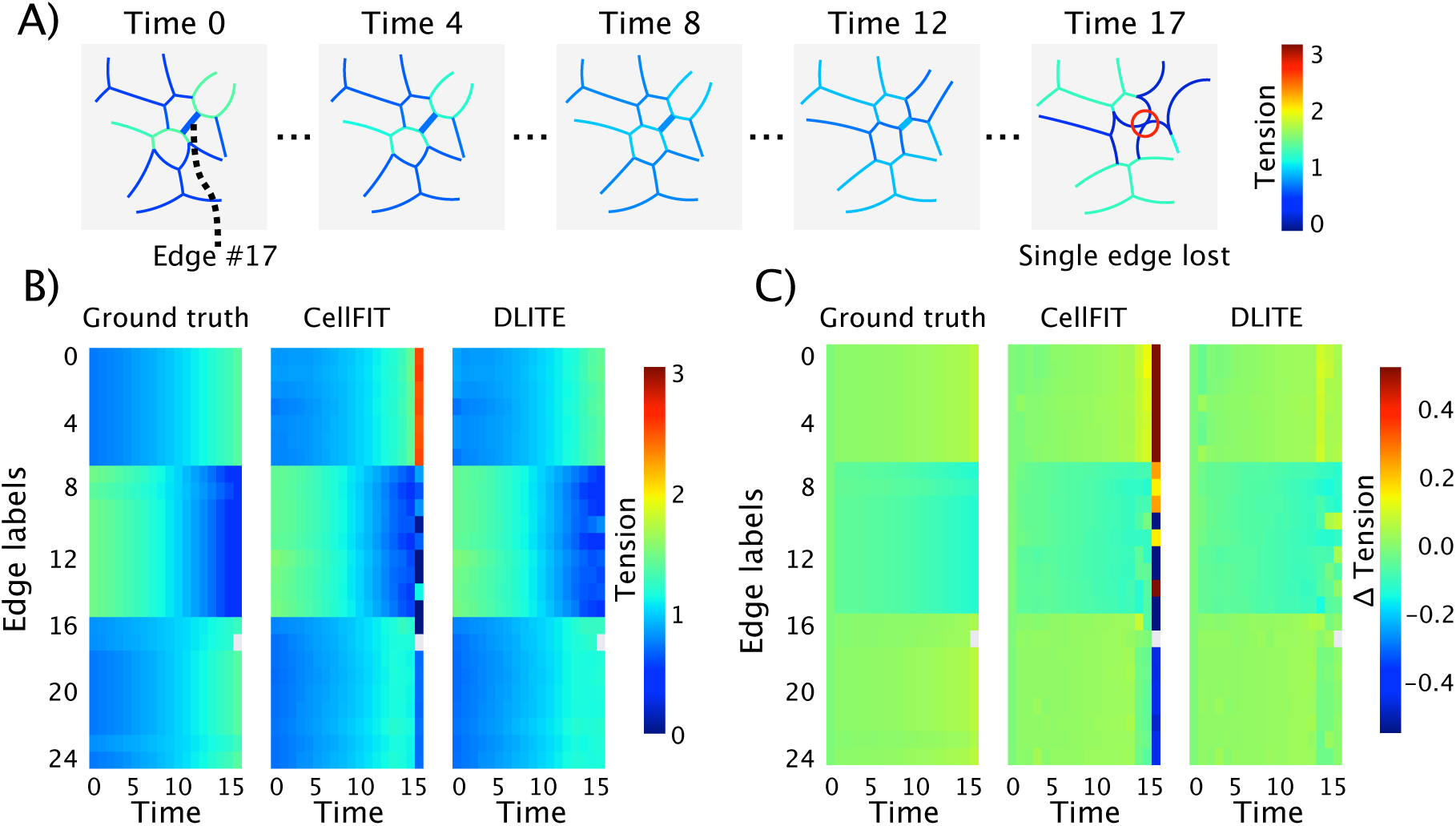
Reduced sensitivity to topological changes in DLITE. (A) Time-series of a synthetic geometry containing 24 edges generated using Surface Evolver where edge label 17 disappears at time 17 (circled in red). (B) Heatmap of dynamic edge tensions for ground truth, CellFIT (r = 0.75), and DLITE (r = 0.98). DLITE shows reduced disruption to tension prediction on topological change and more closely matches the ground truth tension. (C) Heatmap of dynamic change (derivative of tension) in edge tensions for ground truth, CellFIT and DLITE.

**Figure 5:**
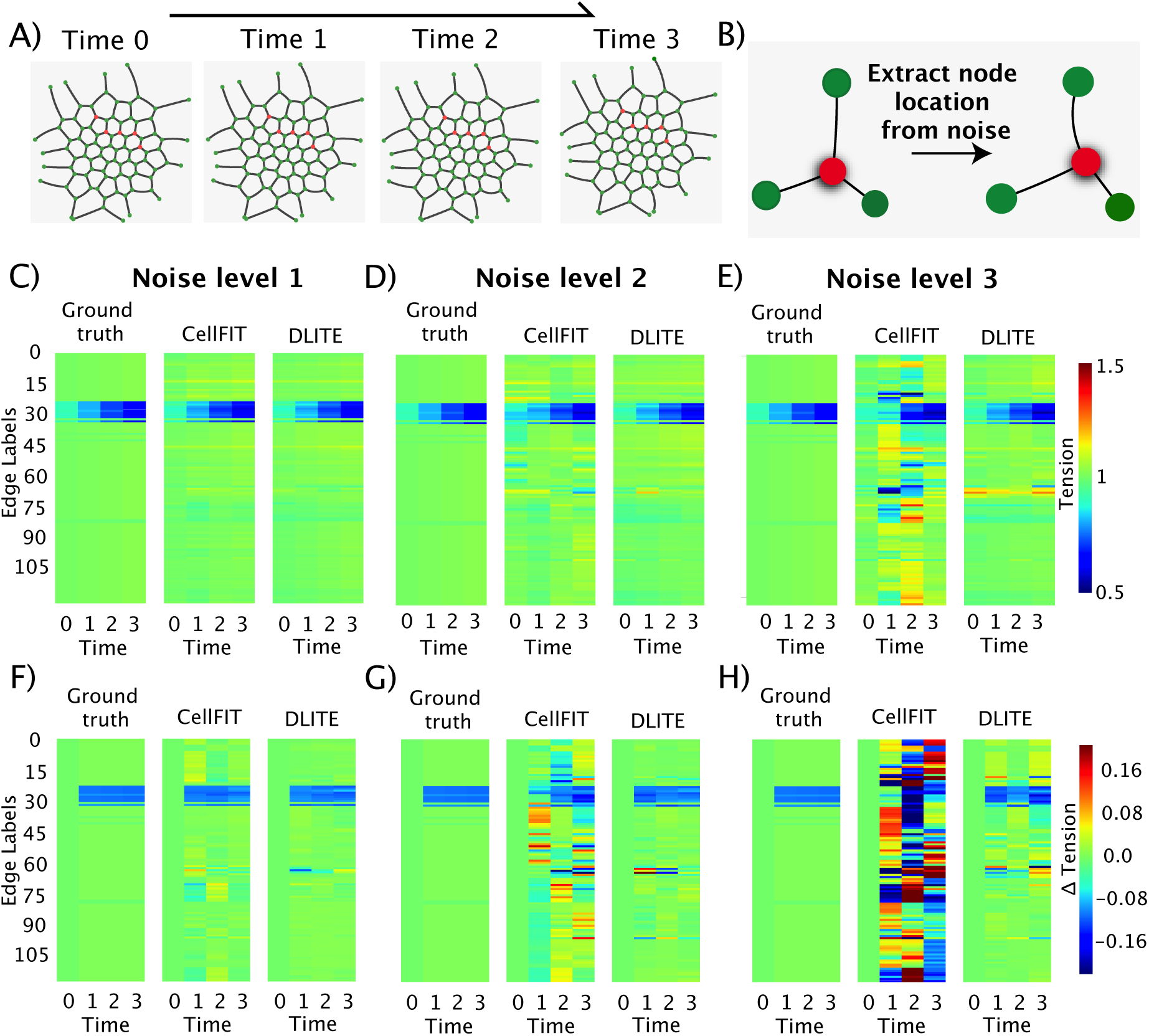
Reduced sensitivity to node location errors in DLITE. Noise levels 1, 2 and 3 correspond to random Gaussian noise added to red node locations, all with a mean 0 and standard deviation 0.1, 0.5 and 1 respectively. (A) Time-series of synthetic colony generated using Surface Evolver. The five nodes subject to perturbation with noise are shown in red. (B) Change in shape of a single triple junction around the red node in the presence of noise. (C, D, E) Heatmap of dynamic edge tensions for ground truth, CellFIT and DLITE at Noise levels 1, 2 and 3 respectively. (F, G, H) Heatmap of dynamic change (derivative of tension) in edge tensions for ground truth, CellFIT and DLITE at Noise levels 1, 2 and 3 respectively.

### Application to ZO-1 tight junctions

Finally, we applied DLITE to experimental images of colonies of hIPS cells. ZO-1 in hIPS cells was tagged at its endogenous locus with mEGFP and visualized using a spinning confocal disk microscope (see SOM for more details). We chose this system because tight junctions (or *zonulae occludentes*) are known to form a selective barrier, regulating paracellular diffusion through the spaces between cells. Injury of tight junctions can impair barrier function, leading to complications in lungs [58, 59], kidneys [60], eyes [61] or the small intestine [62]. The actin cytoskeleton plays an important role in the regulation of this barrier function [63], and is connected to the rest of the tight junction complex through ZO-1 proteins [64, 65]. Recent studies suggest that actin polymerization and transient Rho activation (‘Rho flares’) act to quickly restore barrier function upon localized ZO-1 loss at cell-cell contacts [66]. Mechanical cues from the polymerization and branching of the actin network can lead to reshaping of tight junctions, resulting in varying barrier phenotypes.

Using a skeletonization of segmented GFP images, we predicted the evolution of intercellular forces in three different ZO-1 time series using both DLITE and CellFIT (Fig. 6A, B and C). Since no ground truth is available in this case, we determined the quality of predicted tensions using condition numbers (*κ*) of the tension matrix (Eq. 6) and tension residuals. We note that the relative distribution of tensions range from 0 to 3, such that the average tension is normalized to 1. The time interval between adjacent time points was 3 minutes. The example frames shown in Fig. 6A, B, C are organized as raw GFP (upper), CellFIT predicted tensions (middle) and DLITE predicted tensions (lower). In every frame, we observed at least one kind of digitization error, leading to poor tension matrix condition numbers or tension residuals. Single frame errors such as curvature errors (Fig. 6A - Time 0, *κ* = 69 and Fig. 6B - Time 0, *κ* = 23), connectivity errors (Fig. 6A - Time 10, *κ* = 136 and Fig. 6C - Time 6, *κ* = 10^18^), node location errors (Fig. 6B - Time 5, *κ* = 10^16^ and Fig. 6C - Time 0, *κ* = 10^16^) and time-series specific errors such as new edges (Fig. 6A - Time 5, *κ* = 32.5), missing edges (Fig. 6C - Time 3, *κ* = 31) and topological changes (Fig. 6B - Time 25, *κ* = 46) result in loss of tension stability and errors (Fig. 6A, CellFIT). Despite these digitization errors, DLITE shows increased tension stability (Fig. 6A, DLITE), demonstrating its utility. Heatmaps of dynamic edge tension and change in edge tension (Δtension) for the time-series in Fig. 6 are also shown in Fig. S9. The improved performance of DLITE is predicated on reduced tension residuals at every time point (Fig. S10). Importantly, the reduction in tension residuals is accompanied by a reduced dynamic change in edge tension (Δtension, Fig. S9), indicating a smoothness across time.

**Figure 6:**
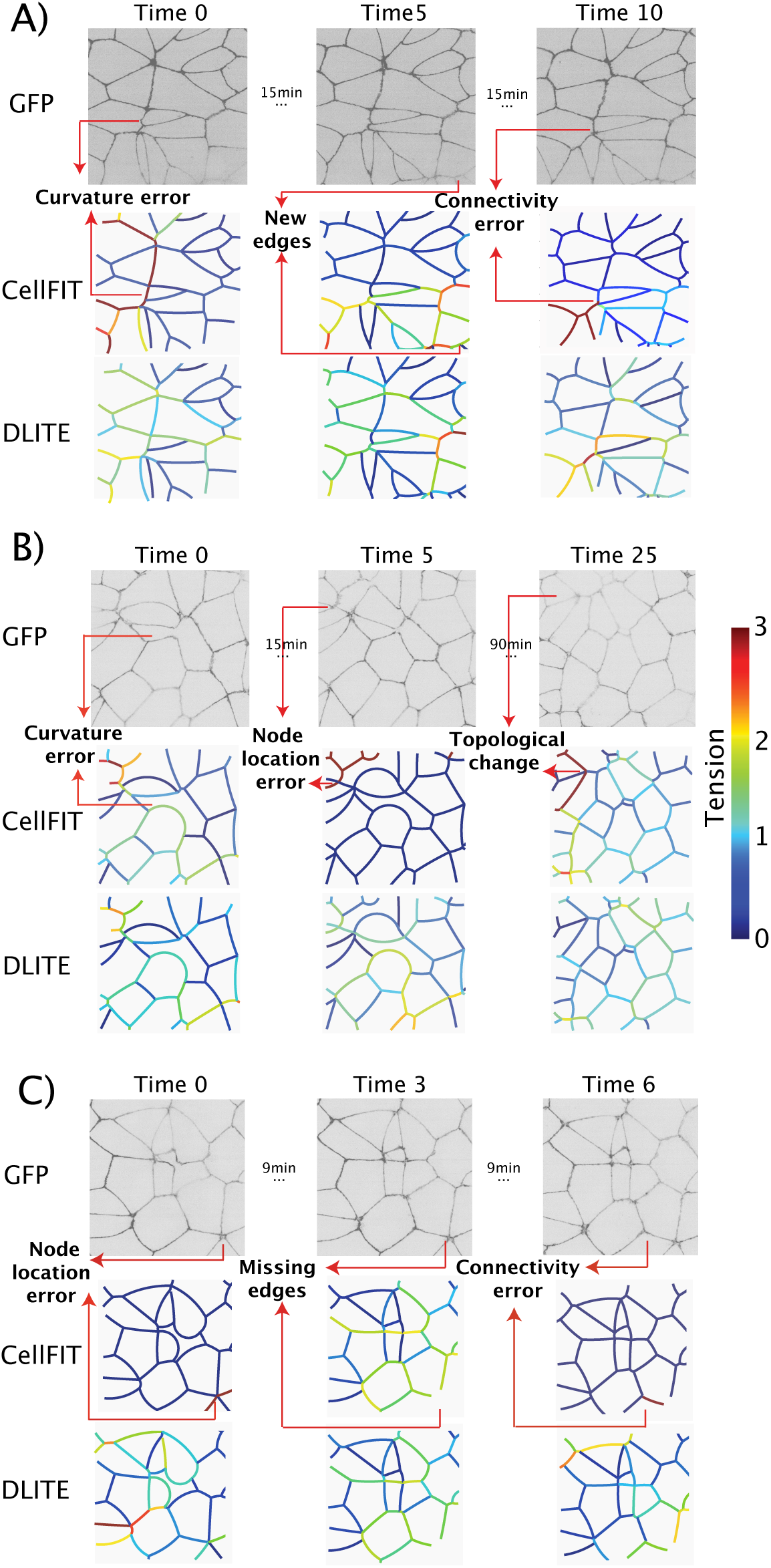
DLITE shows increased tension stability during tension inference in mutliple time-series of ZO-1 labeled hIPS cells. Example frames from three time series are shown in A, B and C and arranged as ZO-1 GFP (upper) and colony edge tensions predicted by CellFIT (middle) and DLITE (lower). Here we use *κ* to denote the condition number of the tension matrix G_*γ*_ (Eq. 6). (A) DLITE shows increased stability to curvature errors (Time 0, *κ* = 69), new edges (Time 5, *κ* = 32.5), connectivity errors (Time 10, *κ* = 136). (B) DLITE shows increased stability to curvature errors (Time 0, *κ* = 23), node location errors (Time 5, *κ* = 10^16^) and topological changes (Time 25, *κ* = 46). (C) DLITE shows increased stability to node location errors (Time 0, *κ* = 10^16^), missing edges (Time 3, *κ* = 31) and connectivity errors (Time 6, *κ* = 10^18^).)

Interestingly, we observed an increase in tension adjacent to a dividing cell immediately after a mitotic event (Fig. 7A, red box) in a time-series of ZO-1 GFP with a single mitotic event at time point 14. This increase in tension post-mitosis was observed using both methods (Fig. 7C, edge labels 3, 10 and 11), but only after the removal of digitization errors during a semi-manual skeletonization process. This step was important to ensure non-poorly scaled CellFIT solutions, such as the ones at time points 2 and 13. As before, both the tension residuals (Fig. 7B) and the dynamic change in tension (Δtension, Fig. 7D, E) were reduced when using DLITE. The reduction in Δtension was determined to be sensitive to the time interval. The standard deviation of Δtension across time was significantly reduced at a time lag of 1 frame (3 minutes), but showed no difference between methods for a time lag of 5 frames (15 minutes).

**Figure 7:**
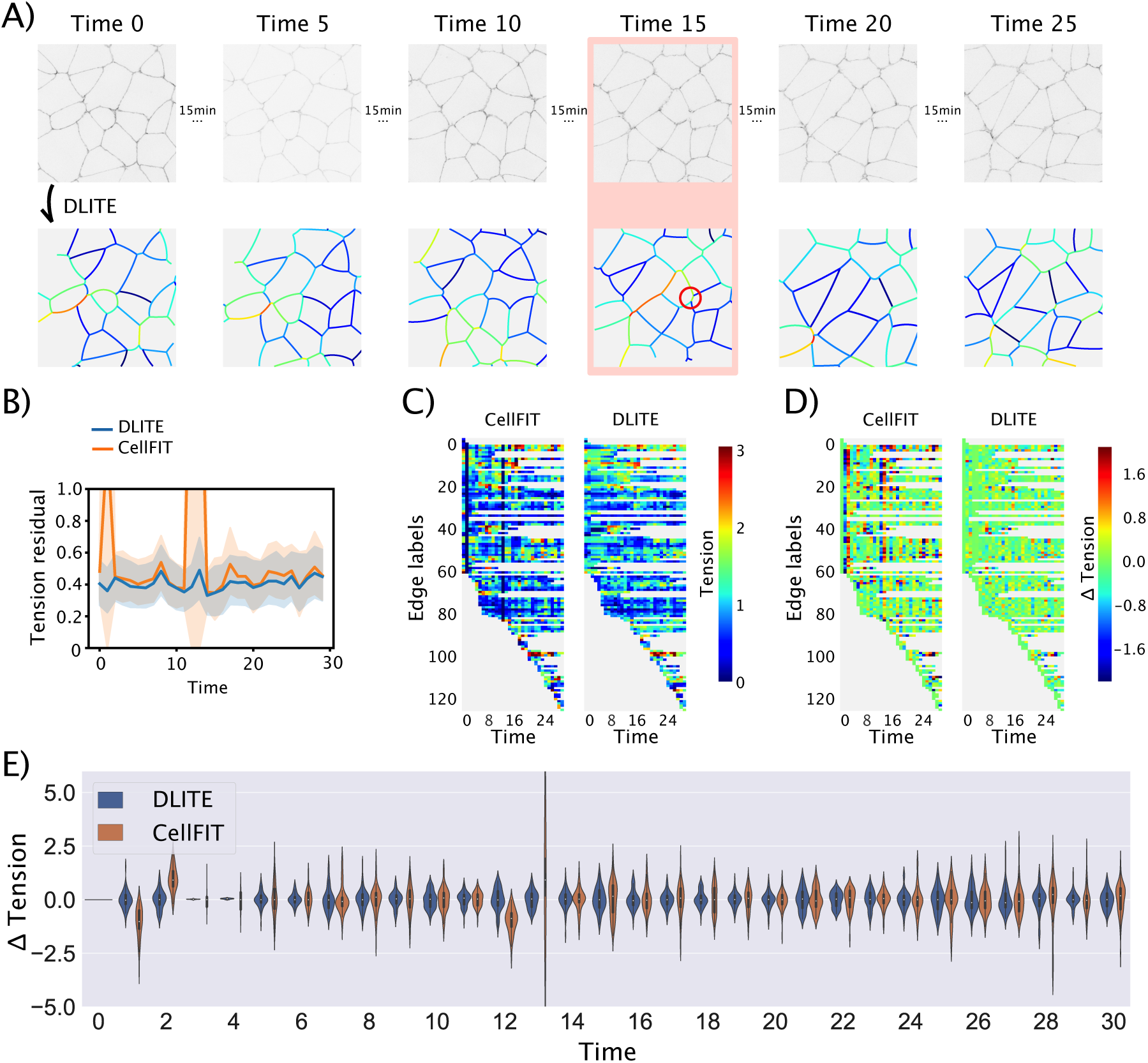
Dynamic cell-cell forces from a time-series of ZO-1 tight junction locations in hIPS cells. DLITE shows reduced fluctuation in tension change, showing more temporally correlated tension predictions. (A) Time-series of ZO-1 GFP images (upper) and dynamics of colony edge tension predicted by DLITE (lower). The following time points are shown: 0, 5, 10, 15, 20 and 25. Time 15 shows an increase in tension along a ridge in the middle of the colony following a mitotic event and the forming of a new edge (circled in red). The time interval between adjacent time points was 3 minutes. (B) Tension residuals at every time point showing an estimate of central tendency and corresponding confidence interval. (C) Heatmap of dynamic edge tensions predicted by CellFIT and DLITE. (D) Heatmap of dynamic change (derivative of tension) in edge tensions predicted by CellFIT and DLITE. (E) Distribution of Δtension (derivative of tension) at every time point for CellFIT and DLITE.

## Discussion

In this study, we have presented a new method, DLITE, which is based on a local optimization of tension residuals to compute dynamic cell-cell forces. We validated the predictive power of DLITE using synthetic geometries generated by Surface Evolver [49] and showed that DLITE performs better than the prior state-of-the-art method CellFIT [34] when applied to time-series data. Importantly, this method incorporates a framework to track nodes, edges, and cells across time.

We demonstrated that DLITE is robust to digitization challenges common in time series data such as poor estimates of edge angle, errors in node location, connectivity errors and topological changes that occur as cells move and encounter different neighbors. Finally, we applied DLITE to estimate edge tensions in multiple time-series of ZO-1 tight junctions and showed improved stability in tension predictions and an increase in tension post mitosis. We indicated that DLITE displays a reduced Δtension compared to CellFIT, indicating greater temporal smoothness. We observed this reduction in three other scenes of the ZO-1 tight junction.

The need for dynamic force-inference tools to understand cell shape and colony rearrangement is driven by their applicability to morphogenic processes from wound healing to germ-band extension to colony reorganization [1, 2, 3, 4]. These processes rely on transient mechanical forces that are ideally detected by the extended non-perturbing observations for which DLITE is designed. Computing the dynamics of cell-cell forces via this computational framework complements experimental advances and enable data-driven estimation of intercellular forces, particularly as biological data sets grow in size.

While useful, DLITE makes assumptions about the system that create limitations. Specifically, DLITE assumes 1) edges are circular arcs, 2) that tension is correlated from timepoint-to-timepoint, and 3) that sufficient computational resources are available. 1) The tensions calculated using DLITE depend on fitting circular arcs to every edge. This approximation breaks down if the edge is not under sufficient tension or the cytoskeleton is strongly perturbing it inhomogeneously across the interface. Under these conditions the inferred tensions will not approach the ground truth. 2) Using local optimization seeded with tensions from the prior time point assumes that the tensions are correlated across these time points. This is evidently true as the inter-frame interval approaches zero and evidently false as the same interval approaches infinity. DLITE’s informed prior decreases in usefulness as we increase this interval or the timescale of our system’s force variance decreases. 3) Finally, as the amount of biological data increases, implementation of DLITe must be optimized for computation speed in large colonies.

DLITE offers comparable tension inference to existing methods when applied to single time points, increased performance when applied across time points, increased stability in the face of segmentation challenges, and increased stability when applied to limited experimental data sets. Future use of DLITE will look at dynamic changes in cell-cell forces in larger data sets of ZO-1 tight junctions, allowing the visualization of cell-cell forces during large scale colony reorganization.

## Acknowledgments

We thank the entire Allen Institute for Cell Science team, who generated and characterized the gene-edited hiPS cell lines and developed image-based assays used in this work. We especially thank the Allen Institute for Cell Science Assay Development team, particularly Irina Mueller for collecting the ZO-1 time series used here and Susanne Rafelski for invaluable discussions and advise. We would like to thank the Allen Institute for Cell Science Animated Cell team and Daniel Toloudis specifically for providing the 3D rendering of ZO-1 cells used in Figure 1. We thank Paul G. Allen, founder of the Allen Institute for Cell Science, for his vision, encouragement and support. We would also like to thank members of the Rangamani lab and Dr. Matthew Akamatsu for comments and feedback. This work was partially supported by ARO W911NF1610411 to P.R.

## Supplementary Online Material

### Data structures

We implemented our code using the standard scientific Python stack. An object/class groups similar constructs together. Here, we defined 4 main objects - nodes, edges, cells and colonies. Nodes are objects with a unique location (*x, y*) and node label. Edges are objects that are connected to two unique nodes with a defined edge curvature, direction and edge label. Cells are objects with a unique cell label that contain a particular list of nodes and edges, where a combination of the contained edges forms a cycle. Colonies are objects comprising a list of cells and stray edges (edges that are not part of any cell). Each class has several other defined properties that were useful for time-series tracking and cell-cell force inference.

### Curve-fitting

We fit a circular arc to a given list of (*x, y*) edge co-ordinates using a least squares fitting routine from the Python module Scipy [40].

### Cell finding algorithm

Given a list of nodes and edges, we loop through every edge to find the two (or fewer) cells of which each edge might be a part. We start from an initial edge and find the closest edge that forms the smallest (or largest) angle with the current edge. We repeated this process by setting the new edge as current edge, until the second node of the current edge is identical to the first node of the initial edge, thus indicating a complete cycle. We validated this algorithm in planar graphs generated using the NetworkX module [67]. In pseudocode, this can be formulated as shown in Algorithm 1.

### Surface Evolver simulations

Synthetic geometries were generated using the Surface Evolver [49], which provides a precise way to make model geometries using soap film physics. An initial surface was first defined in a datafile comprising a list of vertices, edges, facets and bodies, along with any volume or area constraints.

**Algorithm 1.**
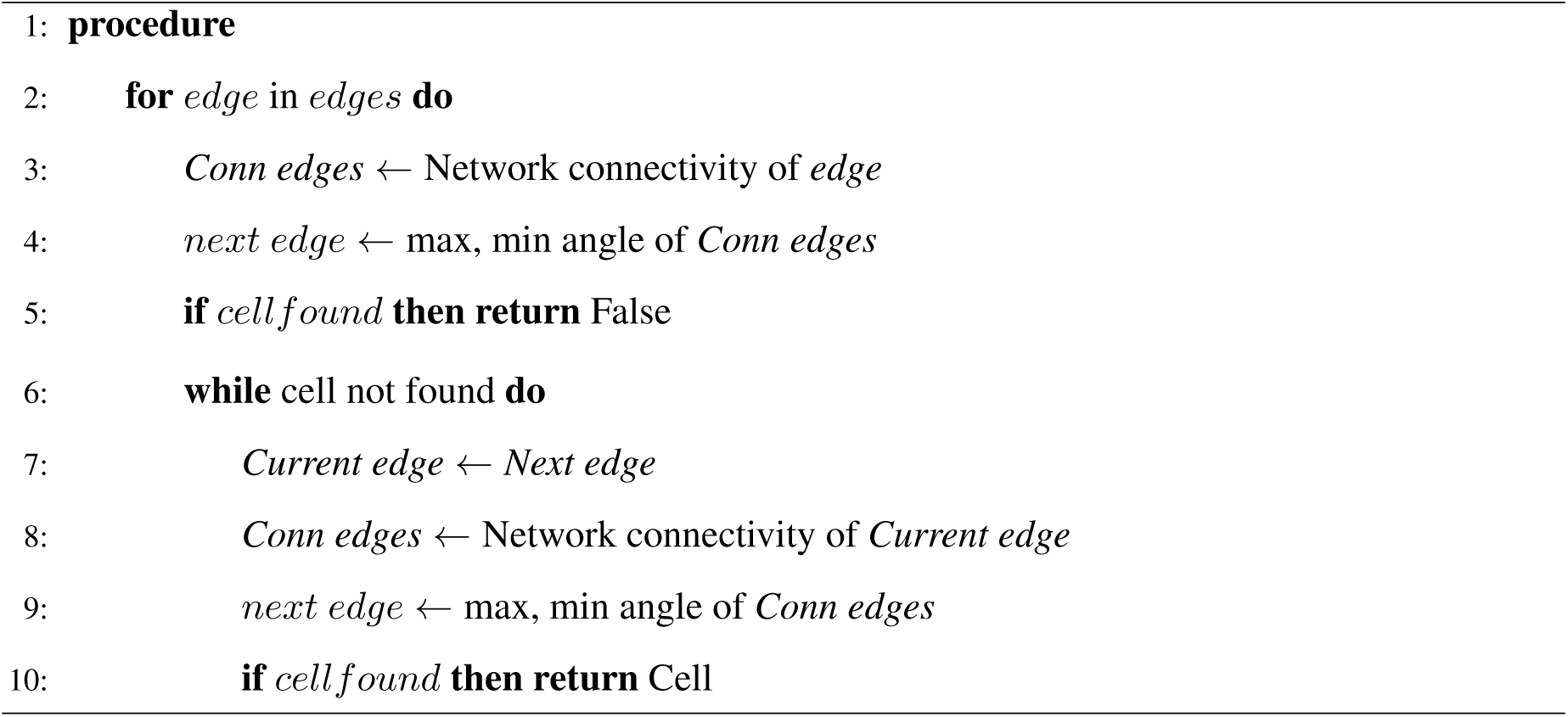
Cell finding algorithm

Since we defined a 2D string model with a space dimension of 2, we enforced area constraints on the facets instead of volume constraints on the bodies. To generate the datafile, we made random Voronoi tessellations. This was followed by Lloyd relaxation to make a more uniform tessellation. Edges were then assigned random tensions. All cells were assigned the same fixed area in a single simulation (typically 5000). Evolver then uses gradient descent to morph the surface to a minimum energy (*W*) configuration. This energy (*W*) was defined as Eq. 5 with the pressure energy enforced as an area constraint. Multiple mesh refinement steps (adjustments of every vertex in the system) were used to ensure a minimum of 10 mesh points along every edge (for better curve-fitting). The evolved geometries were screened for edges that form cell-cell interfaces (as opposed to edges along the boundary that do not form an interface). These edges, along with their corresponding nodes and cells, were stored using our data structures.

In order to generate a time-series, we took a given geometry and perturbed the tension of random edges. Surface Evolver was used to solve for the minima. This was the next frame t + 1. We repeated this tension perturbation procedure uniformly, such that the geometry was smoothly changing its curvature along every edge in correlation with a changing tension (Eq. 5). By stitching together multiple Evolver geometries that progressively showed increasing or decreasing curvature/tension of certain edges, we were able to generate movies of colony rearrangement that was smooth across time, for whom the ground truth tensions were the input to Surface Evolver.

### CellFIT solution

CellFIT [34] evaluates the tension balance as a matrix system defined as

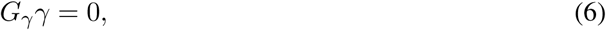

where *γ* is a list of surface tension magnitudes and G_*γ*_ is a matrix of edge tension coefficients (sin’s and cos’s). Since the system of equations is over-determined, this is formulated as a constrained least squares (Karush Kuhn Tucker or KKT) matrix, which can be written as

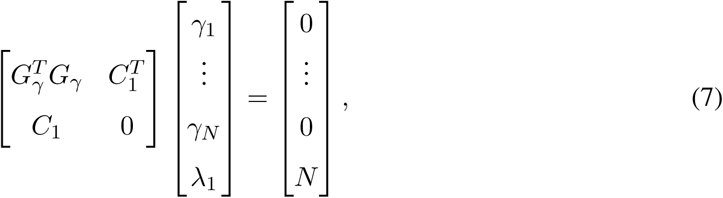

where C_1_ = [1, …, 1], *λ*_1_ is a Lagrange multiplier, and N is the number of edge tensions. This normalizes the average edge tension to 1. Similarly, the pressure balance is evaluated as a matrix system

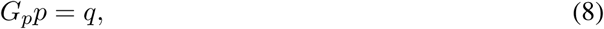

where *p* is a matrix of cell pressures and *q* is a matrix of edge tensions divided by edge curvatures (*t/r*), as per Laplace’s law. This is also formulated as a constrained least squares matrix as

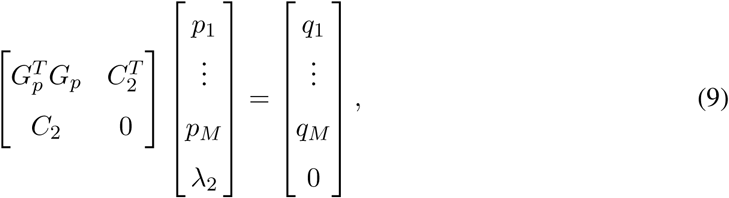

where C_2_ = [1, …, 1], *λ*_2_ is a Lagrange multiplier and M is the number of cell pressures. This normalizes the average edge pressure to 0.

### Force-inference algorithm

In pseudocode, the force-inference algorithm can be formulated as shown in Algorithm 2.

**Algorithm 2.**
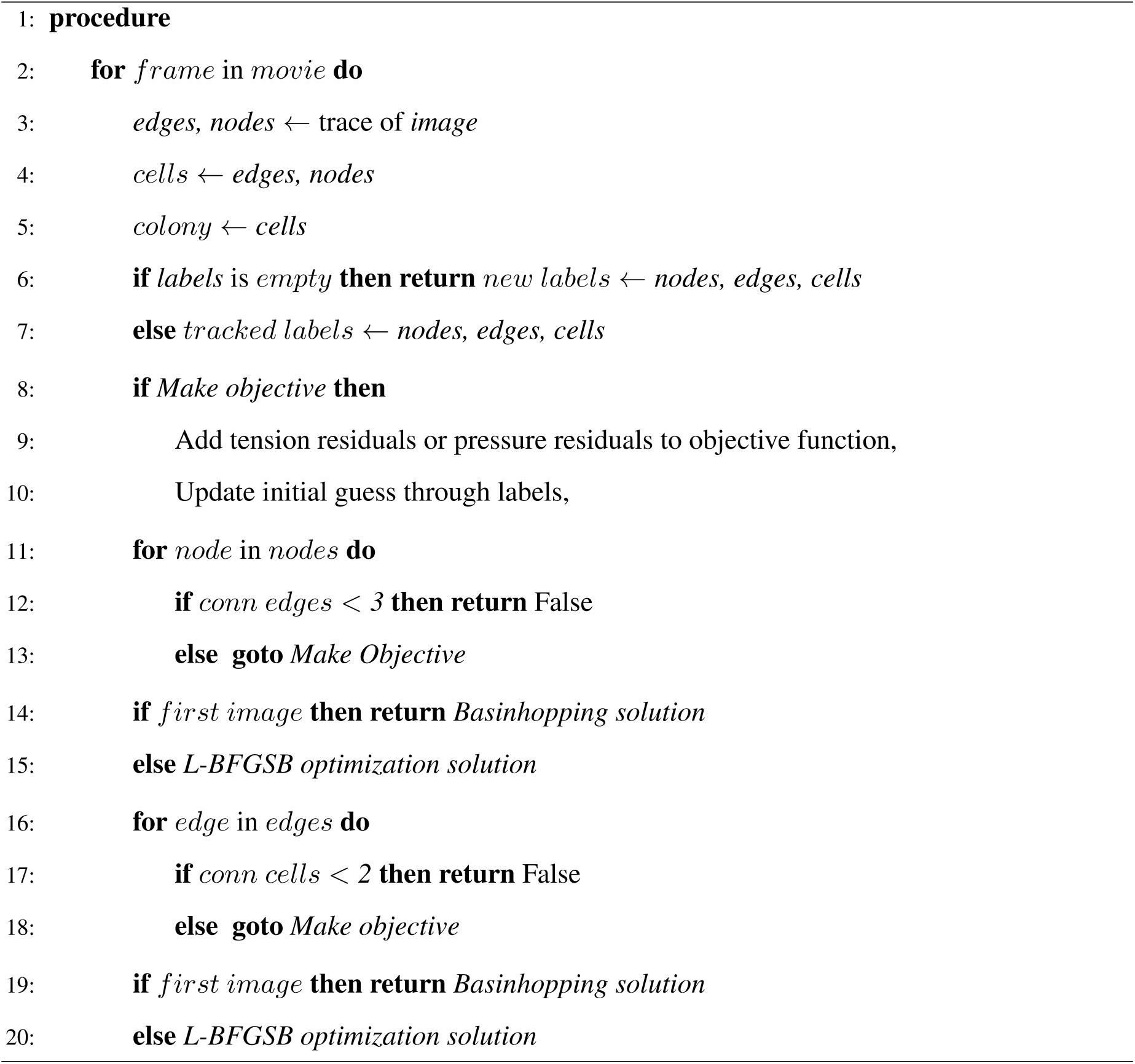
Dynamic cell-cell force inference algorithm

### Cell plating for imaging

Human induced pluripotent stem cells (hiPSCs) were plated on glass-bottom multiwell plates (1.5H glass; Cellvis) coated with phenol red–free GFR Matrigel (Corning) diluted 1:30 in phenol red–free DMEM*/*F12 (Life Technologies). Cells were seeded at a density of 2.5 *×* 10^3^ in 96-well plates and (12.5–18) imaged 3–4 days later. A detailed protocol can be found at the Allen Cell Explorer (Allen Institute for Cell Science, 2017).

### Live-cell imaging

Cells were imaged on a Zeiss spinning-disk microscope with a Zeiss 100*×*/1.25 W C-Apochromat Korr UV Vis IR objective, a CSU-X1 Yokogawa spinning-disk head, and Hamamatsu Orca Flash 4.0 camera. Microscopes were outfitted with a humidified environmental chamber to maintain cells at 37°C with 5 % CO_2_ during imaging. Time-lapse movies were acquired every 3 minutes for 1.5 hours.

**Figure S1:**
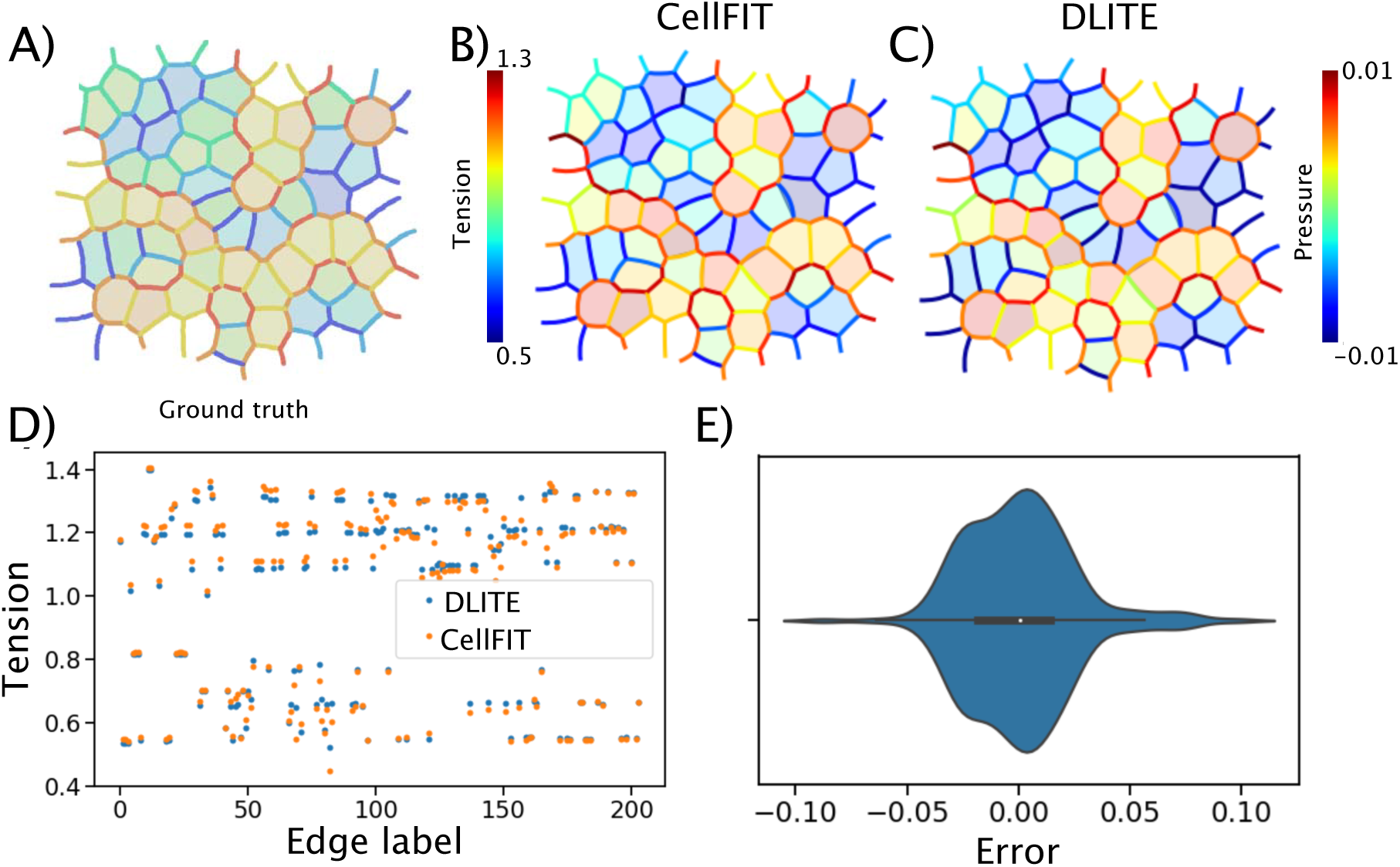
Validation of rewritten CellFIT code. (A) Ground truth geometry used in [34]. (B) Tensions and pressures predicted using CellFIT. (C) Tensions and pressures predicted using DLITE. (D) Tension vs Edge label for CellFIT and DLITE. (E) Error between DLITE tension and CellFIT tension.

**Figure S2:**
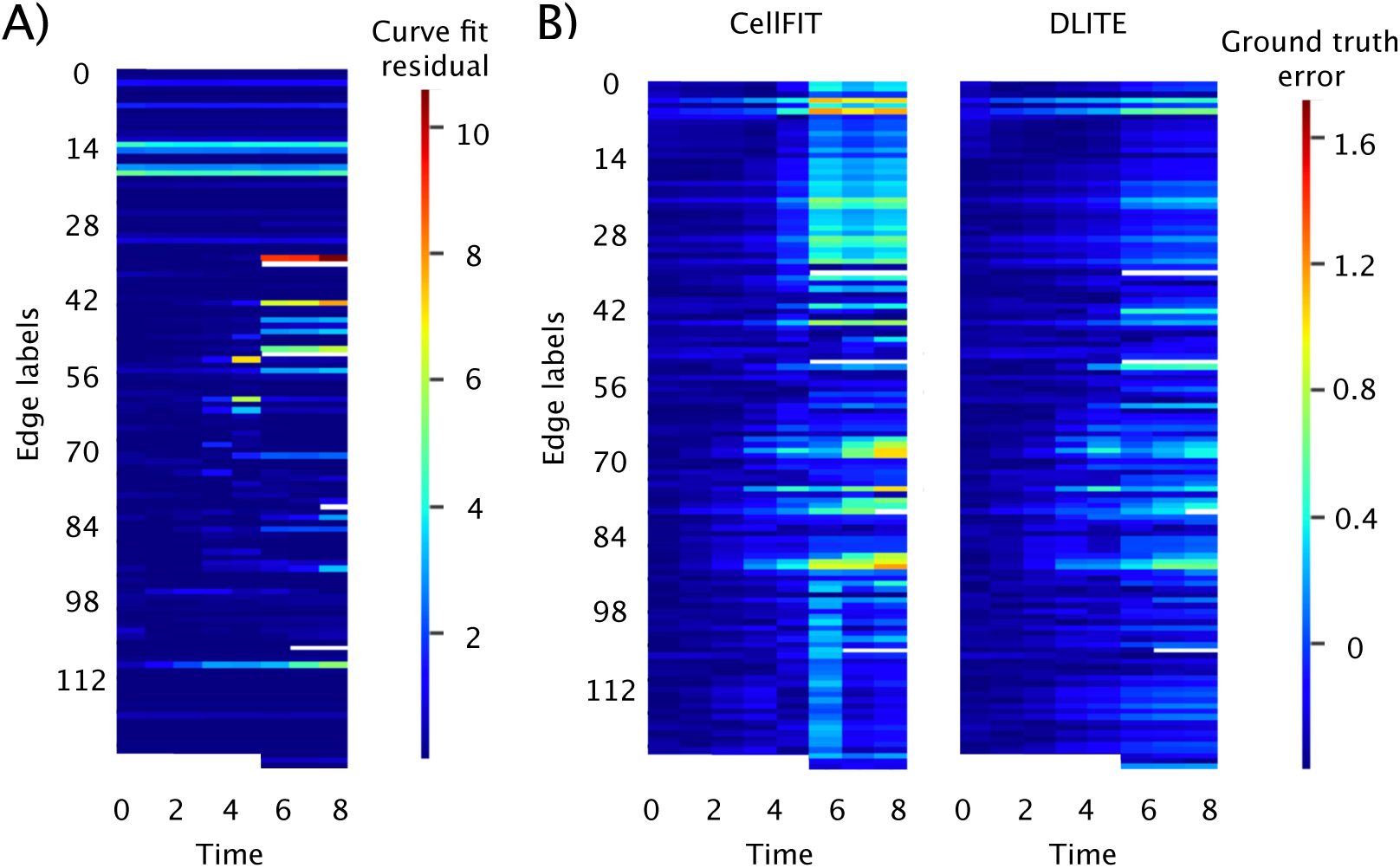
Curve fit residuals and ground truth error for the synthetic colony time-series shown in Fig. 2. (A) Heatmap of curve fit residuals. (B) Heatmap of dynamic ground truth tension errors using CellFIT and DLITE.

**Figure S3:**
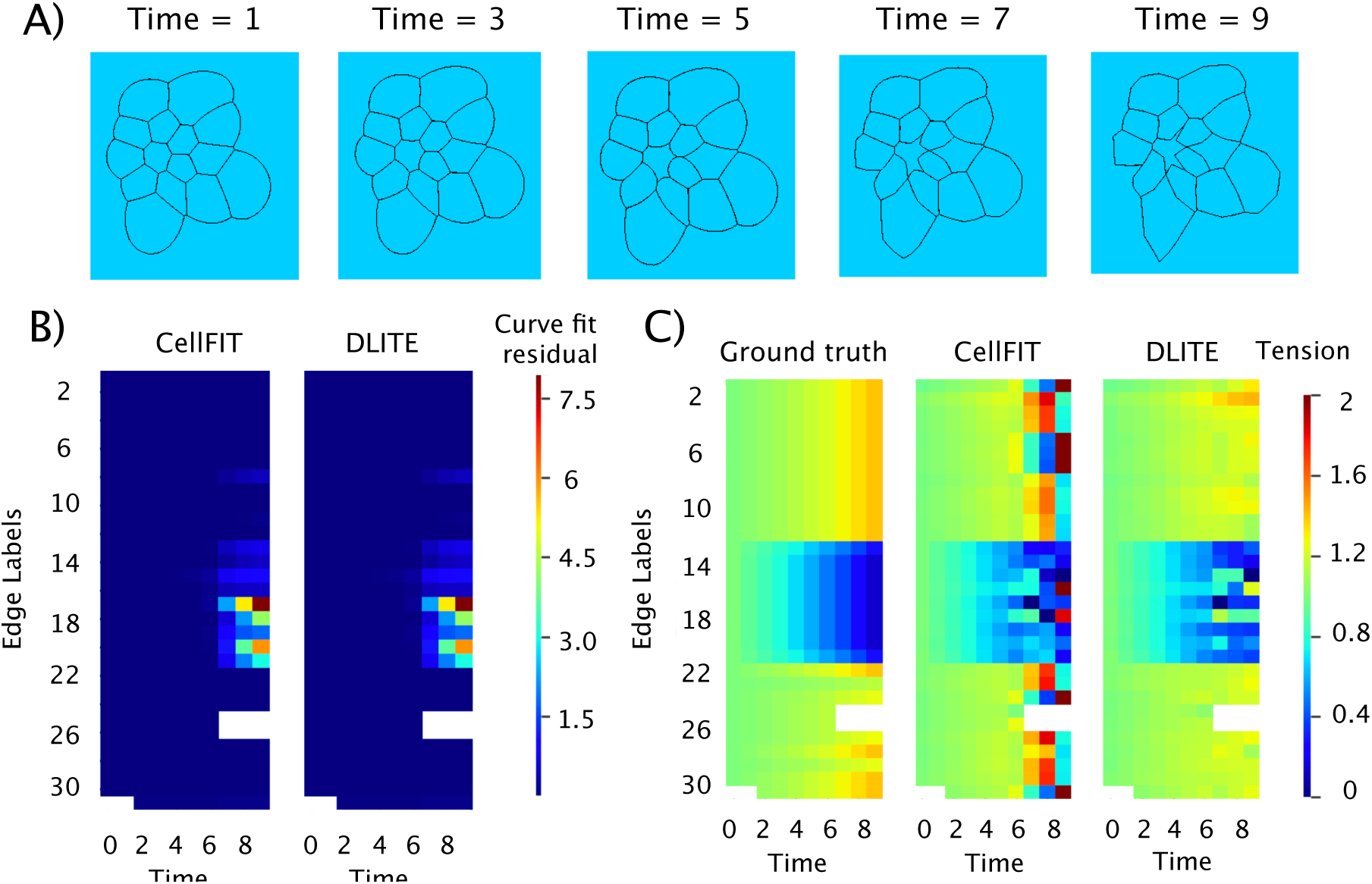
DLITE shows reduced sensitivity to curve fitting errors. (A) Time-series of a synthetic geometry evolved in Surface Evolver with distinctly non circular edges at later time points (Time 7, 9). (B) Heatmap of curve fit residuals. (C) Heatmap of dynamic edge tensions for ground truth, CellFIT and DLITE.

**Figure S4:**
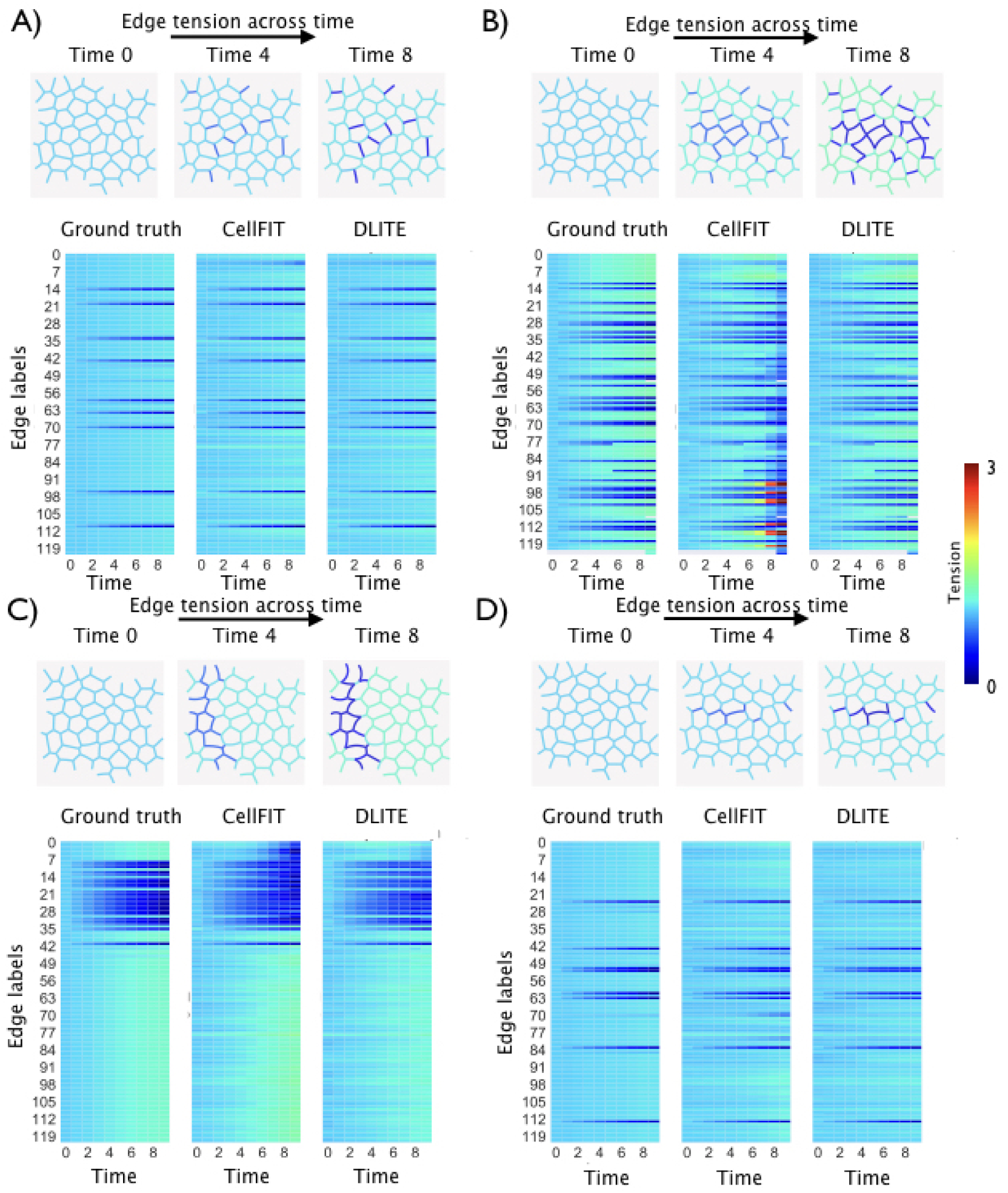
Four example time-series (A-D) of colony rearrangement simulated using four different combinations of decreasing the tension of a few edges and increasing tension of all other edges in the same colony geometry as that in Fig. 2. Average edge tension is normalized to 1 at every time point. Shown - 3 example time points (Time 0, 4 and 8) and heatmaps of dynamic edge tensions for ground truth, CellFIT and DLITE. (A) DLITE - r = 0.97, CellFIT - r = 0.96, (B) DLITE - r = 0.93, CellFIT - r = 0.54, (C) DLITE - r = 0.96, CellFIT - r = 0.86, (D) DLITE - r = 0.94, CellFIT - r = 0.93

**Figure S5:**
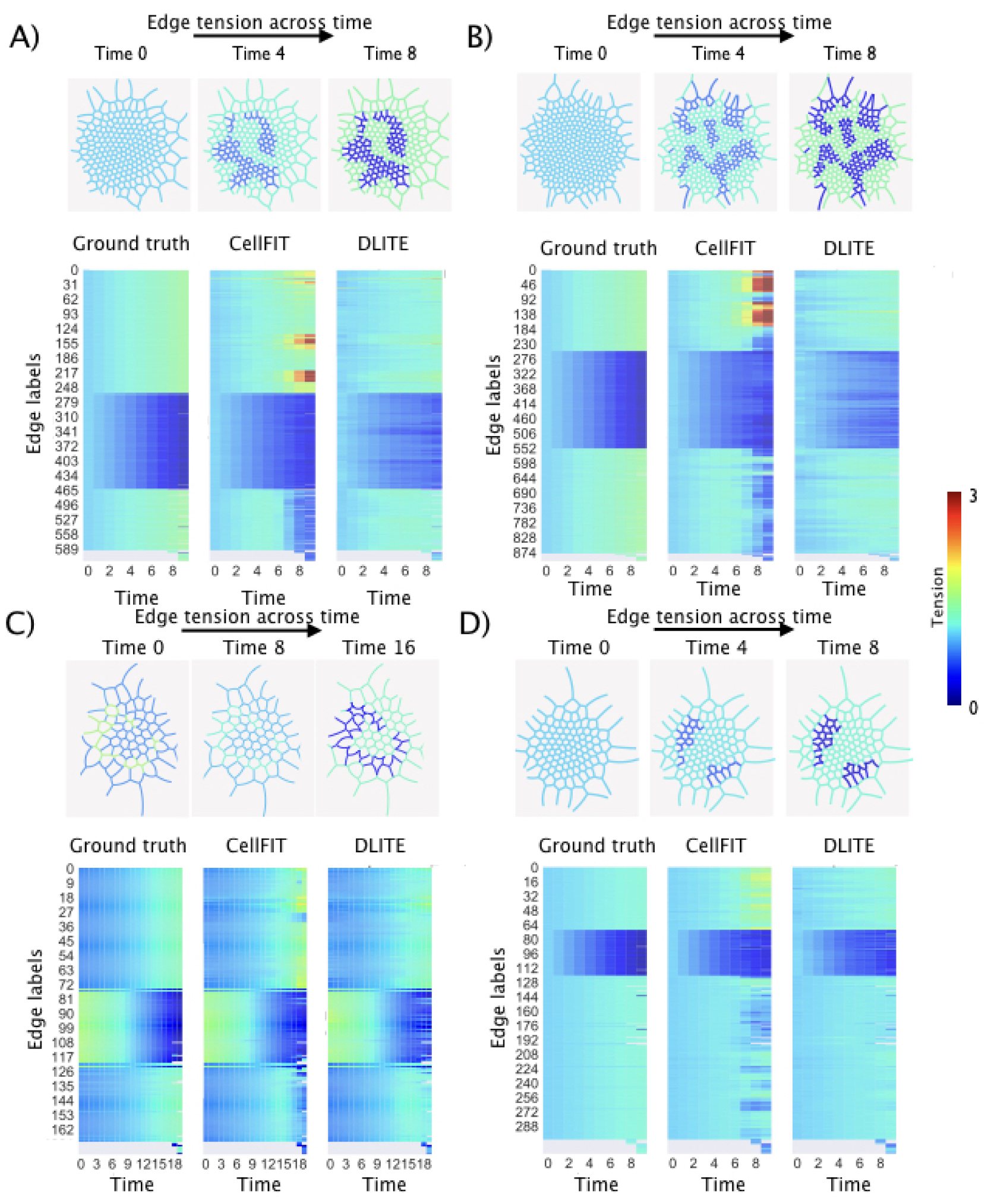
Four example time-series (A-D) of colony rearrangement simulated by decreasing the tension of a few edges and increasing tension of all other edges in four randomly generated colony geometries of different sizes. Average edge tension is normalized to 1 at every time point. Shown - 3 example time points (Time 0, 4 and 8 or Time 0,8 and 16) and heatmaps of dynamic edge tensions for ground truth, CellFIT and DLITE. (A) DLITE - r = 0.94, CellFIT - r = 0.76, (B) DLITE - r = 0.92, CellFIT - r = 0.6, (C) DLITE - r = 0.95, CellFIT - r = 0.9, (D) DLITE - r = 0.96, CellFIT - r = 0.84.

**Figure S6:**
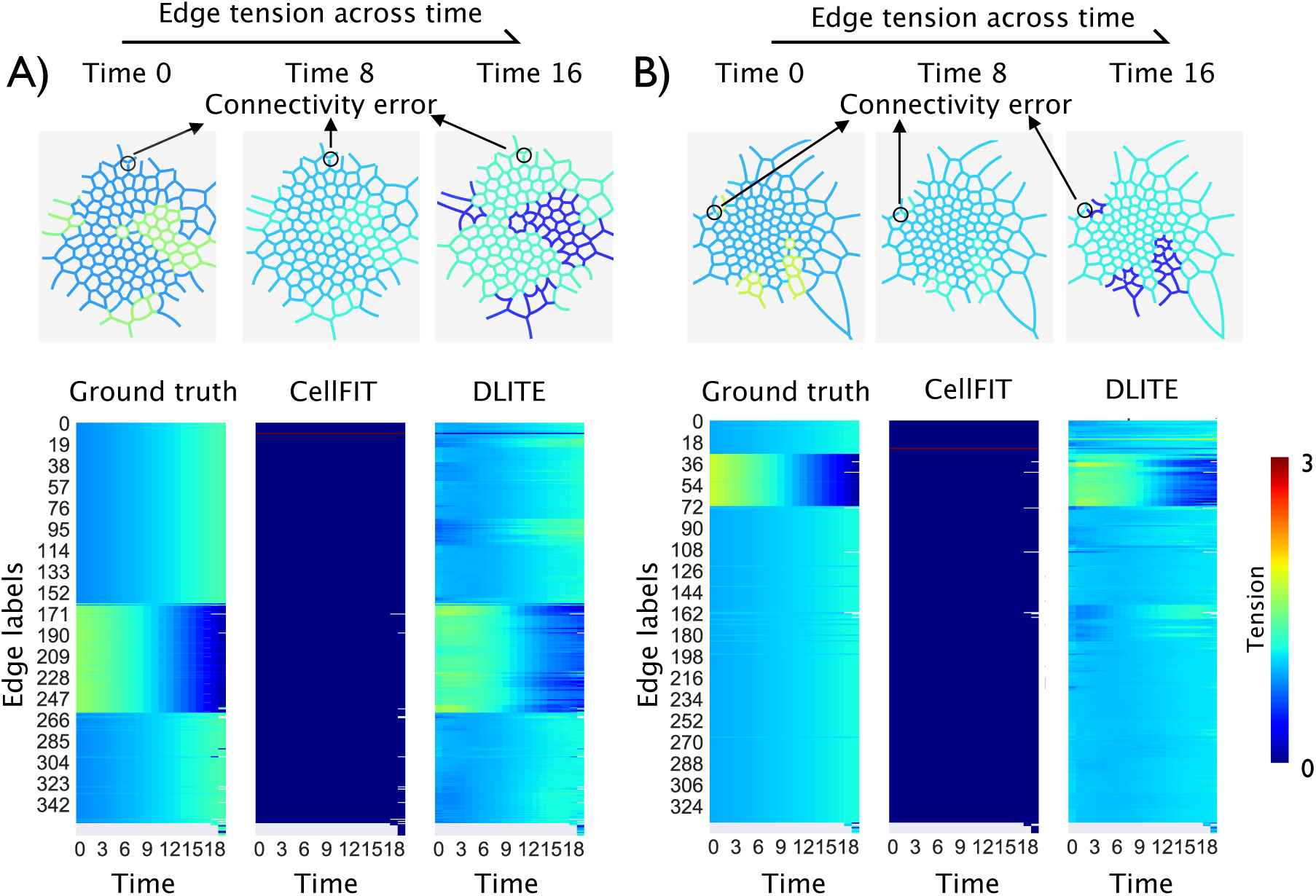
Two example time-series (A, B) of colony rearrangements with single connectivity errors at a node. Average edge tension is normalized to 1 at every time point. Shown - 3 example time points (Time 0, 8 and 16) and heatmaps of dynamic edge tensions for ground truth, CellFIT and DLITE. (A) DLITE - r = 0.88, CellFIT - r = 0.22, (B) DLITE - r = 0.83, CellFIT - r = 0.25.

**Figure S7:**
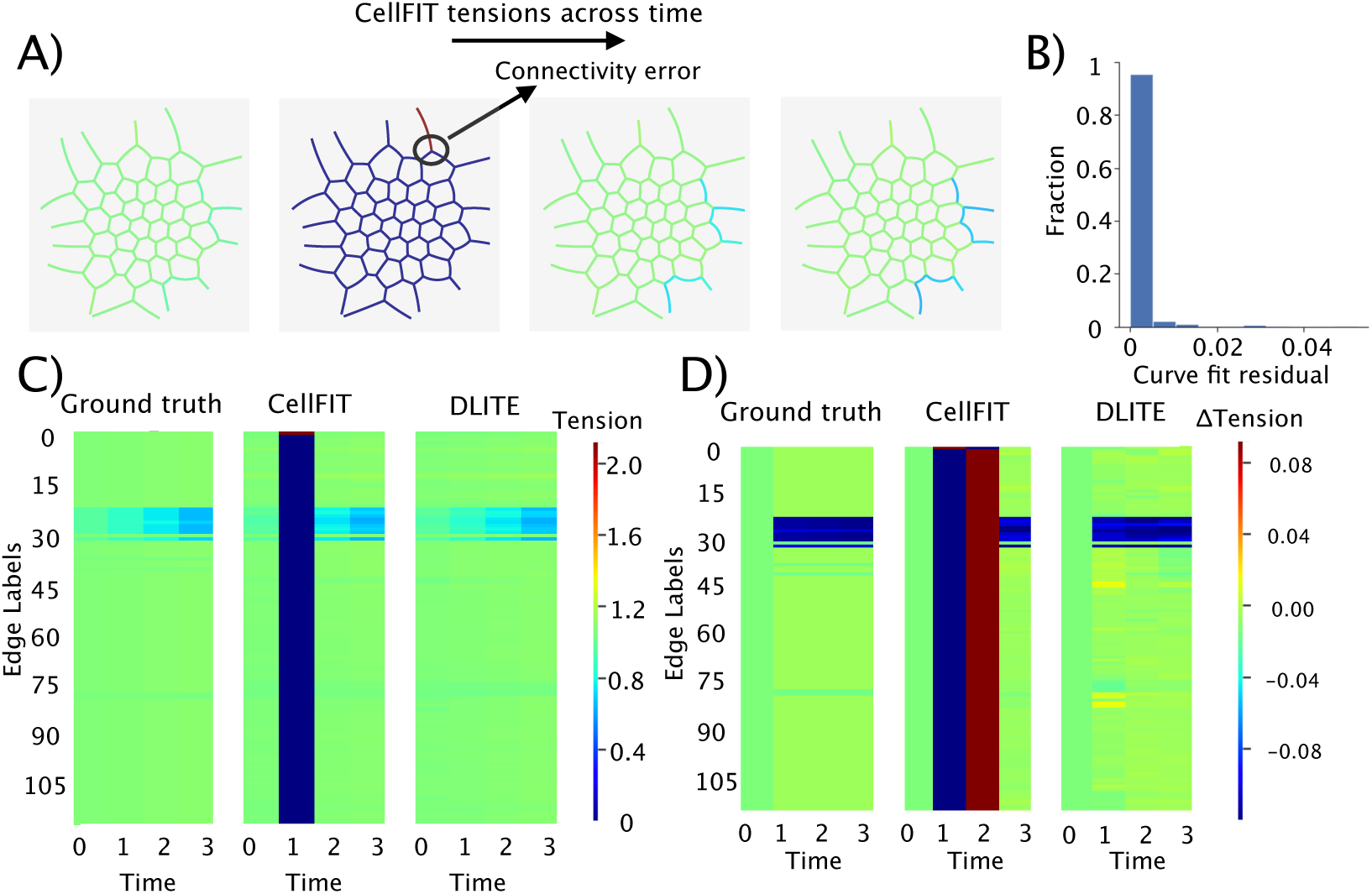
Synthetic colony time-series with a single connectivity error at time point 1. (A) Time-series of colony edge tensions predicted using CellFIT. (B) Histogram of curve fit residuals at all time points. (C) Heatmap of dynamic edge tensions for ground truth, CellFIT and DLITE. (D) Heatmap of Δtension (derivative of tension) for ground truth, CellFIT and DLITE.

**Figure S8:**
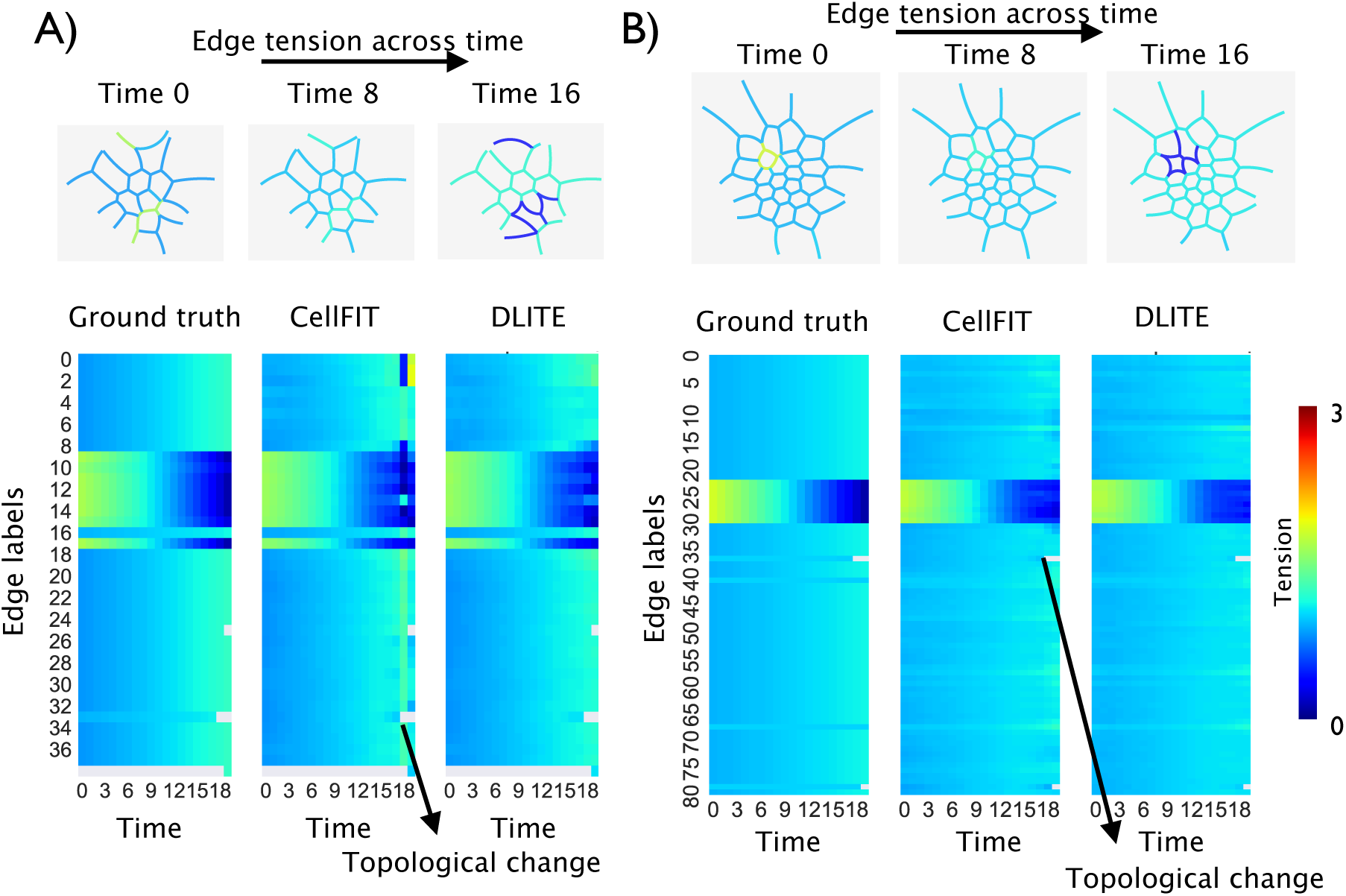
Two example time-series (A, B) of colony rearrangements with single topological changes (shrinkage of cell-cell junctions) at a node. Average edge tension is normalized to 1 at every time point. Shown - 3 example time points (Time 0, 8 and 16) and heatmaps of dynamic edge tensions for ground truth, CellFIT and DLITE. (A) DLITE - r = 0.97, CellFIT - r = 0.91, (B) DLITE - r = 0.975, CellFIT - r = 0.974.

**Figure S9:**
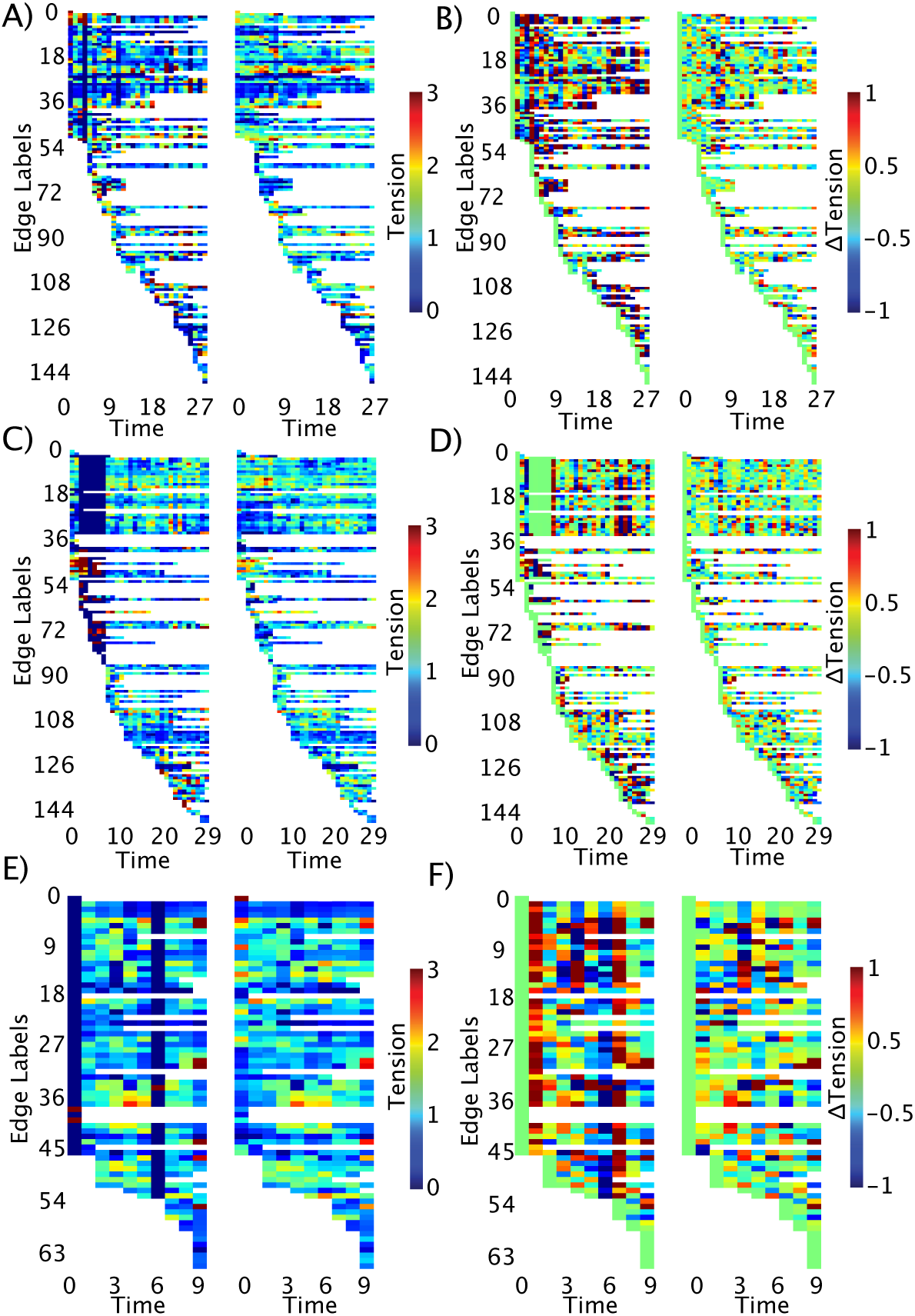
Heatmaps of dynamic edge tension (A, C, E) and dynamic change (derivative of tension) in edge tension (B, D, F) for the ZO-1 time-series shown in Fig. 6A, 6B and 6C respectively.

**Figure S10:**
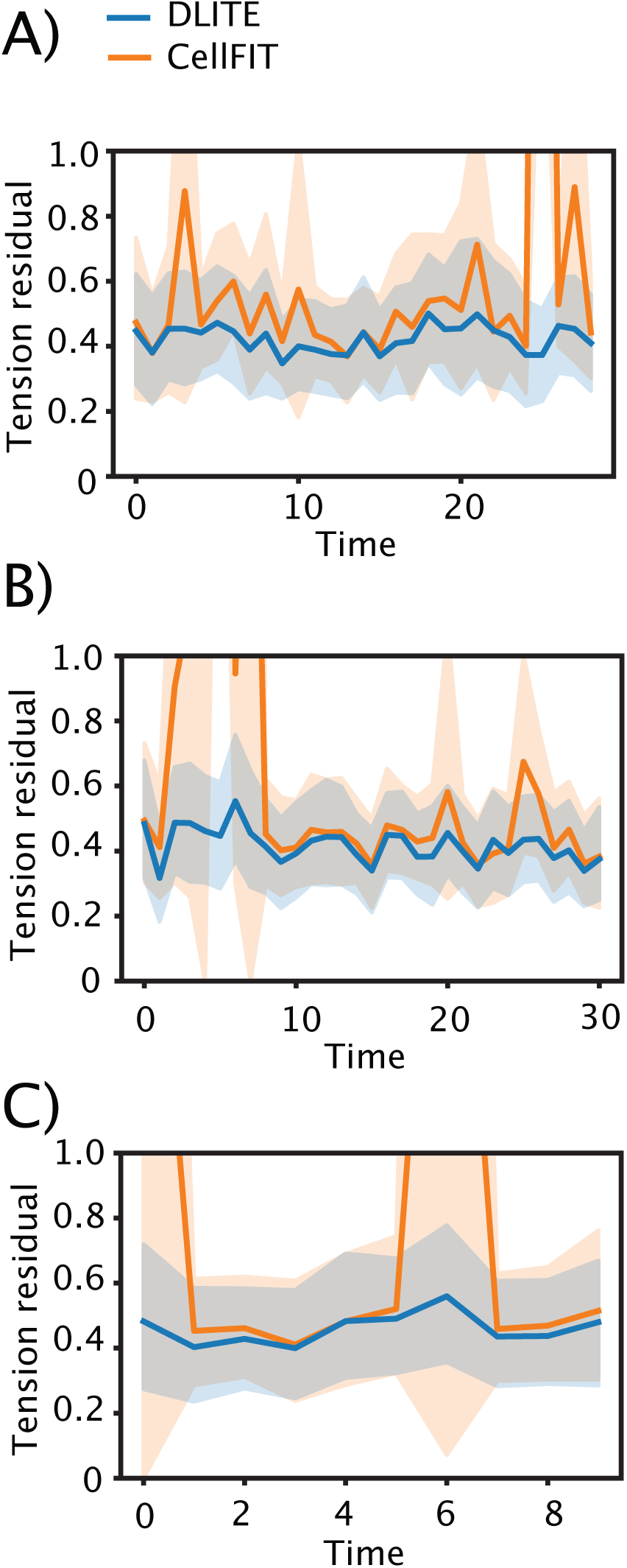
Lineplots (A, B, C) of tension residuals showing an estimate of the central tendency and a confidence interval for that estimate for the ZO-1 time-series shown in Fig. 6A, 6B and 6C respectively.

